# Cell-cell communication through FGF4 generates and maintains robust proportions of differentiated cell types in embryonic stem cells

**DOI:** 10.1101/2020.02.14.949701

**Authors:** Dhruv Raina, Azra Bahadori, Angel Stanoev, Michelle Protzek, Aneta Koseska, Christian Schröter

## Abstract

During embryonic development and tissue homeostasis, reproducible proportions of differentiated cell types are specified from populations of multipotent precursor cells. Molecular mechanisms that enable both robust cell type proportioning despite variable initial conditions in the precursor cells, as well as the re-establishment of these proportions upon perturbations in a developing tissue remain to be characterized. Here we report that the differentiation of robust proportions of epiblast-like and primitive endoderm-like cells in mouse embryonic stem cell cultures emerges at the population level through cell-cell communication via a short-range FGF4 signal. We characterize the molecular and dynamical properties of the communication mechanism, and show how it controls both robust cell type proportioning from a wide range of experimentally controlled initial conditions, as well as the autonomous re-establishment of these proportions following the isolation of one cell type. The generation and maintenance of reproducible proportions of discrete cell types is a new function for FGF signaling that may operate in a range of developing tissues.

## Introduction

The differentiation of specialized cell types from populations of multipotent precursor cells is the basis of embryonic development and tissue homeostasis in the adult. Developing tissues generally produce and maintain a standard end result consisting of defined proportions of differentiated cell types despite biological noise and perturbations, a behavior termed canalization (Waddington, 1942). Molecular mechanisms that contribute to canalized development by controlling the proportions of differentiated cell types remain to be characterized.

Mammalian preimplantation development is a prime example for developmental canalization. The size of the three lineages, trophoectoderm (TE), epiblast (Epi), and primitive endoderm (PrE) is remarkably constant between mouse preimplantation embryos (Saiz et al., 2016). Furthermore, mammalian embryos can regulate the proportions of the three lineages following splitting, fusing, or the addition of embryonic stem cells (ESCs), such that the embryo is capable of post-implantation development (Bedzhov et al., 2014; Bradley et al., 1984; Gardner, 1968; Martinez Arias et al., 2013; Tarkowski, 1961; 1959). The differentiation of Epi and PrE cells from inner cell mass (ICM) precursor cells is controlled by transcription factors such as NANOG and GATA6, which mark and specify Epi and PrE cells, respectively. These factors are initially co-expressed in ICM cells, and become mutually exclusive as cells differentiate (Chazaud et al., 2006; Plusa et al., 2008; Simon et al., 2018). In addition, PrE differentiation requires fibroblast growth factor (FGF)/ extracellular regulated kinase (ERK) signaling (Chazaud et al., 2006; Kang et al., 2017; 2012; Krawchuk et al., 2013; Molotkov et al., 2017). The expression dynamics of lineage-specific transcription factors, together with extensive analysis of genetic mutants and pharmacological interventions in the embryo have led to models in which mutually repressive interactions between transcriptional regulators together with heterogeneous FGF/ERK signaling allocate individual ICM cells to one of the two lineages (Yamanaka et al., 2010; Bessonnard et al., 2014; Chickarmane and Peterson, 2008; De Caluwé et al., 2019; De Mot et al., 2016). More recently, FGF signaling has been shown to underlie the regulation of Epi and PrE-lineage sizes following the addition or ablation of cells, suggesting that it orchestrates cell differentiation at the population level (Saiz et al., 2020).

Population-level mechanisms for cell differentiation are an attractive solution to the problem of developmental canalization, as it has been shown theoretically that populations of communicating cells can re-establish specific cell type proportions following perturbations (Stanoev et al., 2019). This theory furthermore predicts that the differentiation outcomes are robust towards initial conditions, since the heterogeneous cell identities are generated and maintained on the level of the communicating population, rather than being specified intrinsically in each cell. Identifying and experimentally verifying a molecular mechanism that leads to such emergent phenomena however requires an experimental system in which both the initial conditions of the precursor population as well as cell-cell communication can be precisely controlled.

Here we approach this challenge using the differentiation of Epi- and PrE-like cells from ESCs upon the transient expression of exogenous GATA factors as a model system. PrE-like differentiation in ESCs requires above-threshold levels of GATA factor expression and ERK activity (Fig. 1A, (Schröter et al., 2015)), which supports the concept that mutually repressive interactions between NANOG and GATA factors together with FGF/ERK signalling control differentiation in single cells.

**Figure 1.**
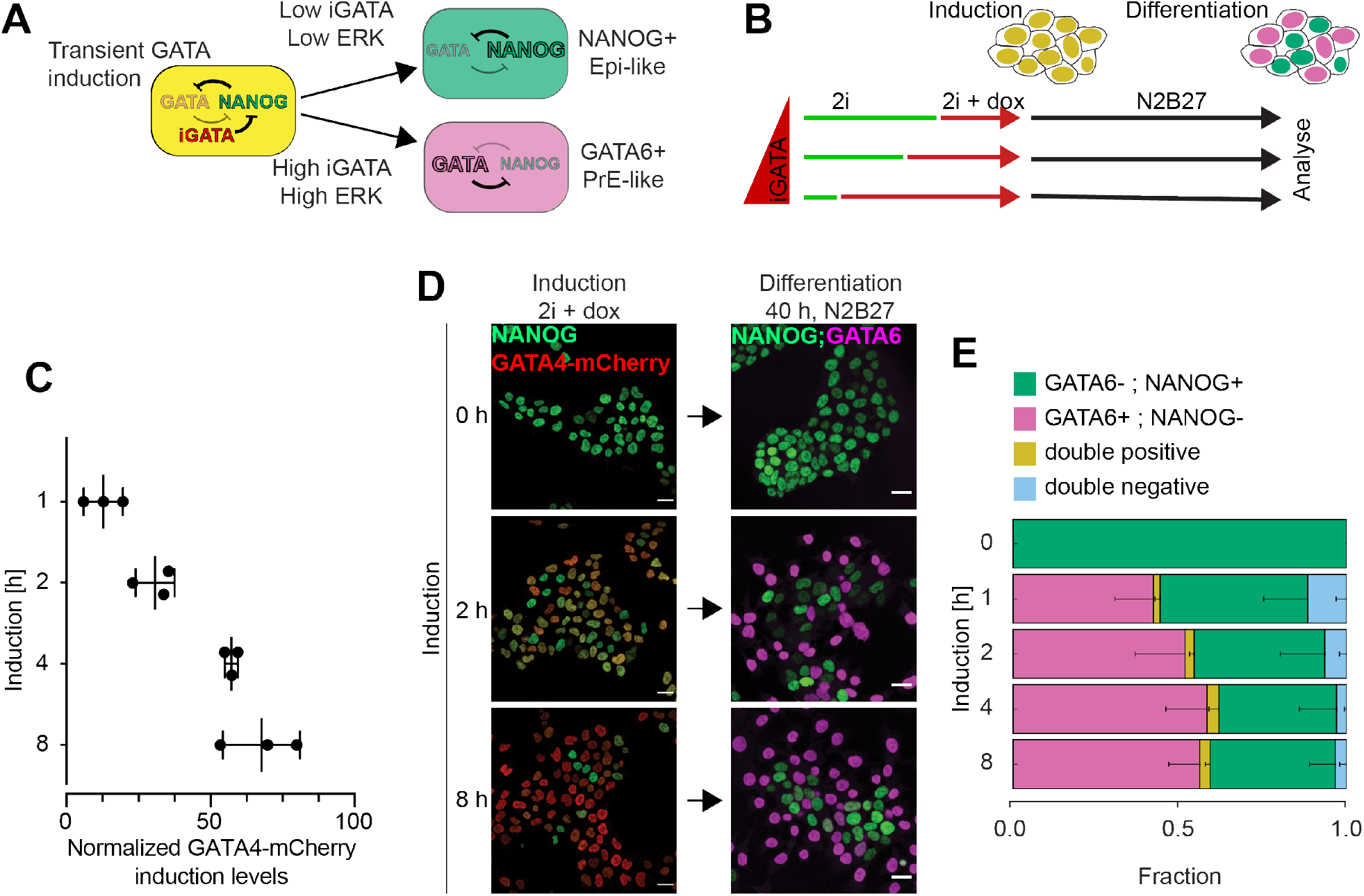
Proportions of differentiated cell types are independent from GATA4-mCherry induction levels. **A** Schematic representation of cell differentiation following transient doxycycline-controlled expression of inducible GATA factors (iGATA). For sufficiently high inducible GATA levels and ERK activity, ESCs undergo PrE-like differentiation marked by endogenous GATA6 expression (magenta, bottom). Otherwise they adopt an Epi-like identity marked by NANOG expression (green, top). **B** Scheme of experimental protocol for titrating inducible GATA4-mCherry expression levels by doxycycline addition to individual samples at different time points. All samples were switched simultaneously to N2B27 medium, such that the total time from seeding to analysis was held constant. **C** GATA4-mCherry expression levels for different durations of doxycycline induction in 2i + LIF medium measured by flow cytometry, normalized to the non-induced control. Individual data points show mean fluorescence intensities from at least 20.000 cells in an individual experiment, bars indicate mean ± SD across three independent experiments. **D** Left: immunostaining for NANOG (green) and GATA4-mCherry (red) in inducible cell lines immediately after the end of a doxycycline pulse of the indicated durations. Right: Immunostaining for NANOG (green) and GATA6 (magenta) in cells treated with doxycycline for the indicated durations, followed by 40 h of differentiation in N2B27 medium. Cells without doxycycline induction have been continuously maintained in 2i medium. **E** Average cell type proportions from N = 4 independent experiments; fraction of GATA6+, NANOG-cells in magenta, GATA6-, NANOG+ cells in green, double positive cells in yellow, and double negative cells in blue. Error bars: 95% confidence interval (CI).

We exploit the ability to experimentally control GATA expression levels in this system to demonstrate that robust proportions of Epi- and PrE-like cells differentiate even when starting from a wide range of initial conditions. Mutant analysis identifies recursive cell-cell communication via local FGF4 signaling as the minimal molecular mechanism for this emergent property of cell type proportioning. These experimental observations can be described as a dynamical property of an inhomogeneous steady state (IHSS), a population-based solution that is characterized with mutually exclusive gene expression patterns (Stanoev et al., 2019). We experimentally confirm a central prediction of this theory by demonstrating that cell-cell communication allows populations to re-establish a mixture of Epi- and PrE-like cells from isolated PrE-like cells. Our work thus provides direct experimental evidence that local cell-cell communication via a secreted growth factor can contribute to robust and reproducible global proportions of differentiated cell types across a wide range of initial conditions.

## Results

### Differentiation of robust proportions of Epi- and PrE-like cells in ESC cultures

To study mechanisms that control the proportions of PrE-like and Epi-like cells differentiating in ESC populations, we integrated a doxycycline-inducible GATA4-mCherry construct via piggybac transgenesis to generate ESC lines that allowed sampling a wide range of GATA4-mCherry expression levels. We induced the transgene while cells were cultured in chemically defined minimal N2B27 medium supplemented with the MEK inhibitor PD0325901 (PD03), the GSK3 inhibitor CHIR99021, and the cytokine LIF (2i + LIF medium, (Ying et al., 2008)) to maintain pluripotency. Differentiation was initiated by concomitantly removing doxycycline and switching to N2B27 medium lacking inhibitors an}d cytokines. We first analyzed how the proportions of Epi- and PrE-like cells evolved over time in a selected clonal line, using the expression of the endogenous NANOG and GATA6 proteins as markers for the two cell types. Mutually exclusive expression of NANOG and GATA6 could already be observed as early as 16 h after the initiation of differentiation. From 40 h onwards, the initially large proportion of GATA6+; NANOG+ double positive cells was strongly reduced, and two cell populations were clearly separated in the NANOG-GATA6 expression space (Figures S1A, S1B). One population encompassed most of the GATA6+; NANOG-(PrE-like) cells. Their proportion increased until 40 h and slightly declined thereafter. The second population encompassed both GATA6-;NANOG+ (Epi-like) cells, whose proportion remained stable throughout the time course, as well as double-negative cells whose proportion increased continuously (Figures S1A, S1B). The clear separation between the two populations suggests that the increase in the proportion of double negative cells at the expense of GATA6+; NANOG-PrE-like cells beyond 40 h is mostly fueled by the downregulation of NANOG expression in the GATA6-negative cell population, combined with a slower proliferation of the GATA6-positive population, rather than by the reversion of PrE-like into double negative cells. Together, these observations therefore suggest that stable proportions of Epi- and PrE-like cells are established at 40 h after the initiation of differentiation.

To determine how expression levels of the inducible GATA4-mCherry protein affect cell type proportions, we titrated GATA4-mCherry expression by staggering the initiation of doxycycline induction in time, while keeping the duration of the entire experiment constant (Figure 1B). GATA4-mCherry levels differed by more than 5-fold between 1 h and 8 h of doxycycline induction in 2i + LIF (Figures 1C and 1D), and were paralleled by a concentration-dependent downregulation of NANOG expression (Figures 1D and S2A). After 40 h of differentiation, both GATA6+;NANOG-PrE-like and GATA6-;NANOG+ Epi-like cells were observed (Figures 1D and S2A). Quantitative immunofluorescence revealed that the proportions of PrE-like and Epi-like cells fell within a narrow range for the different induction times. While the proportion of PrE-like cells slightly increased from 43.6 ± 12.1% for 1 h to 59.7 ± 11.9% and 57.2 ± 8.7% for 4 h and 8 h of induction, respectively, the proportion of Epi-cells was stable for different induction times (minimum 26.9 ± 6.9% for 2 h induction, maximum 30.8 ± 3.7% for 8 h induction) (Figures 1E and S2B). Thus, a wide range of inducible GATA4-mCherry expression levels leads to similar proportions of differentiated cell types. To rule out that the robust proportioning reflected pre-specification of cell types or a limited differentiation potential of a subset of cells, we added recombinant FGF4 during the differentiation phase to boost ERK activity and lower the GATA4-mCherry threshold required for PrE-like differentiation. The presence of FGF4 strongly increased the proportion of PrE-like cells at the expense of Epi-like cells compared to differentiation in N2B27 alone, and reached a maximum of 93.5 ± 2.1% for 8 h induction (Figure S2C).

The observation of robust cell type proportioning was corroborated in four clonal cell lines with independent integrations of the GATA4-mCherry transgene, which displayed a more than 8-fold difference in GATA4-mCherry expression levels following 8 h of doxycycline induction (Figures S3A and S3B). In three out of the four clones, similar proportions of both Epi- and PrE-like cells differentiated upon doxycycline removal and culture in N2B27. The one clone in which fewer PrE-like and more Epi-like cells differentiated had the lowest GATA4-mCherry induction levels (Figures S3C and S3D). In the presence of recombinant FGF4 during the differentiation phase, the fraction of PrE-like cells consistently increased compared to differentiation in N2B27 alone, and mirrored clonal differences in GATA4-mCherry induction levels. In the clonal line with the highest GATA4-mCherry levels, the proportion of PrE-like cells reached a maximum of 98.8 ± 2.0% upon addition of exogenous FGF4 (Figure S3D), indicating that almost all cells have PrE-like differentiation potential following sufficiently strong GATA4-mCherry expression. Taken together, these results suggest that the robust proportioning of cell types is established at the population level through cell-cell signaling.

### Differentiating ESCs communicate via a local FGF4 signal

To identify candidate mechanisms for cell-cell communication that could underlie this robust cell type proportioning we focused on FGF4, since FGF/ERK signaling is required for PrE differentiation both in ESCs and in the embryo (Kang et al., 2012; Krawchuk et al., 2013; Kunath et al., 2007; Schröter et al., 2015), and paracrine FGF4 is the main activator of ERK in ESCs (Kunath et al., 2007).

To investigate how GATA4-mCherry induction levels affect FGF4 signaling in the cell population, a *Sprouty4* ^*H2B-Venus*^ (*Spry4* ^*H2B-Venus*^) transcriptional reporter construct that we have previously established as a quantitative readout for long-term FGF4 signaling (Morgani et al., 2018) was integrated in the inducible cell lines. Longer doxycycline induction times corresponding to higher GATA4-mCherry expression levels resulted in reduced mean fluorescence of the *Spry4* reporter after 24 h of differentiation (Figure 2A). This indicates that GATA4-mCherry expression negatively regulates FGF4 signaling during cell type specification. *In situ* mRNA staining (Choi et al., 2018) revealed that this regulation occurred at the transcriptional level. While *Fgf4* mRNA was strongly expressed in most cells before induction in 2i medium (Figure 2B, left), *Fgf4* staining was strongly reduced after 8 h of GATA4-mCherry induction, and confined to cells with low GATA4-mCherry expression levels (arrowheads in Figure 2B, middle). After 40 h of differentiation in N2B27, *Fgf4* expression was mutually exclusive with *Gata6* mRNA expression (Figure 2B, right). This demonstrates that *Fgf4* mRNA expression is negatively correlated with GATA factor expression and PrE-like differentiation in ES cells.

**Figure 2.**
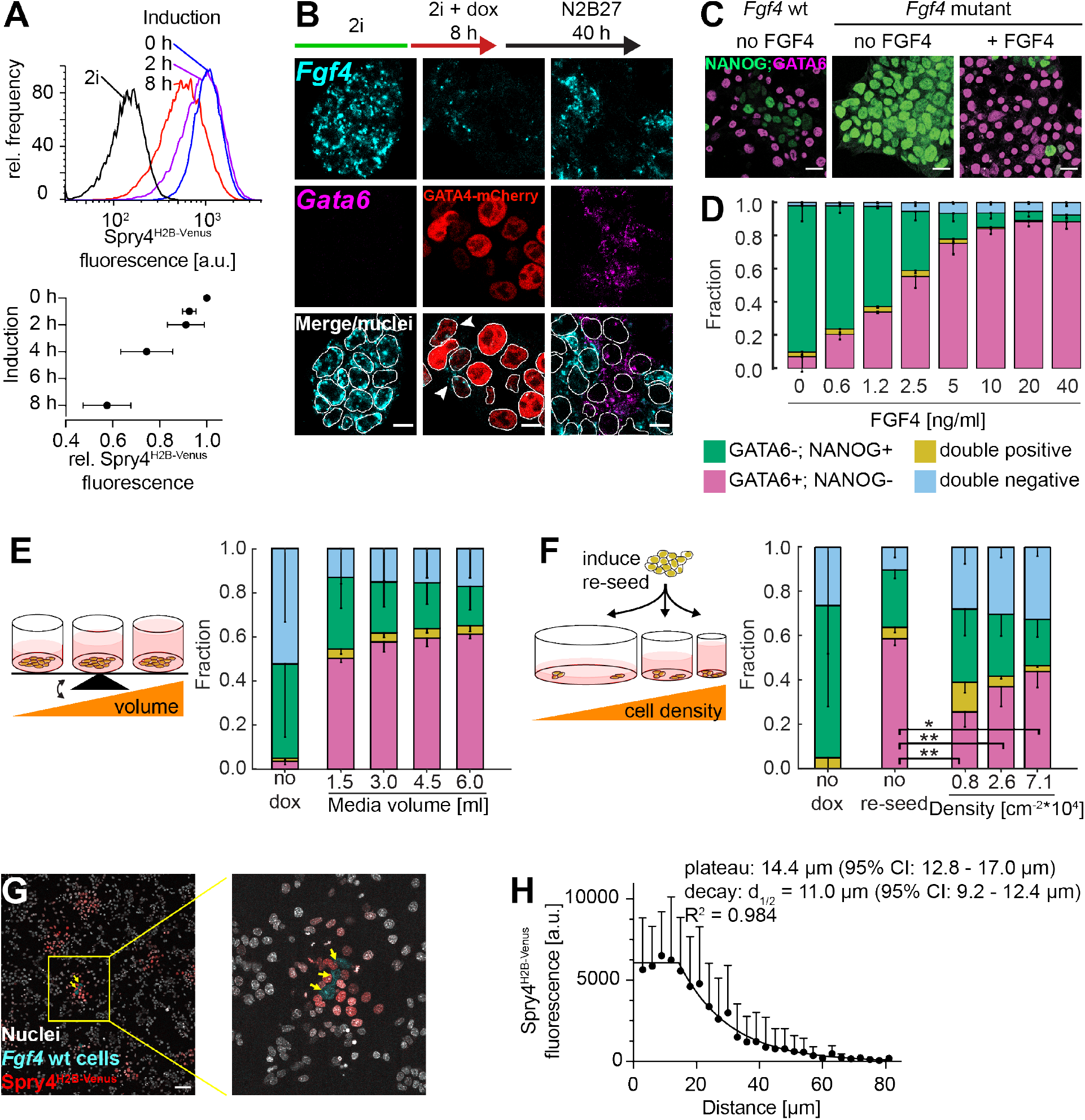
Differentiating ESCs communicate via a local FGF4 signal. **A** Top: Flow cytometry histograms depicting expression of a Spry4^H2B-Venus^ reporter integrated into GATA4-mCherry inducible cells, quantified after 24 h of differentiation in N2B27 medium following the indicated durations of doxycycline induction. Black line indicates reporter expression in cells maintained in 2i medium. Bottom: Mean ± SD of reporter expression from N = 4 independent experiments, normalized to fluorescence levels of cells transferred to N2B27 without doxycycline induction. **B** Fgf4 (top, cyan) and Gata6 (middle, magenta) mRNA staining by in situ hybridization chain reaction at indicated times during the differentiation time-course. At the end of the induction period, GATA4-mCherry protein (red) instead of Gata6 mRNA is detected (middle row, middle panel). Bottom row: Merged images with outlines of nuclei indicated by thin white lines. Scale bar, 10 µm. **C** Immunostaining for GATA6 (magenta) and NANOG (green) in wild type (left) and Fgf4 mutant cells differentiated for 40 h in N2B27 without or with (right) 10 ng/ml FGF4 after an 8 h doxycycline pulse. Scale bars, 20 µm. **D** Average proportions of cell types in Fgf4 mutant cells induced with doxycycline for 8 h, followed by differentiation in N2B27 in the presence of the indicated concentrations of FGF4. GATA6 and NANOG expression were detected by immunostaining and measured by flow cytometry (see Figure S4). N = 4; fraction of GATA6+; NANOG-cells in magenta, GATA6-; NANOG+ cells - green, double positive cells - yellow, and double negative cells - blue. Error bars: 95% CI. **E** Left: Schematic representation of experimental setup to test effects of media volume on cell type proportions. Right: Average cell type proportions determined by flow cytometry, N = 3. **F** Left: Schematic representation of experimental setup to test effects of cell density on cell type proportions. Right: Average cell type proportions determined by flow cytometry, N = 4. ∗∗ and ∗ indicate p < 0.005 and 0.05, respectively (one-way ANOVA). Color code and error bars in **E, F** same as in **D. G** Single labeled Fgf4 wild-type cells (cyan, yellow arrowheads) seeded on a layer of Fgf4 mutant Spry4^H2B-Venus^ transcriptional reporter cells. Nuclei are labeled by siR-Hoechst (white), H2B-Venus fluorescence in red. Scale bar, 100 µm. **H** Quantitative analysis of FGF4 signaling range. Data points show mean ± SD of background-subtracted H2B-Venus fluorescence intensities in nuclei of Spry4^H2B-Venus^ reporter cells quantified in distance bins of 3 µm width relative to the center of mass of Fgf4 wild type cells. Data from N = 9 independent signaling centers. The fluorescence decay length was estimated by fitting a plateau followed by one-phase exponential decay to the data (black line).

We next tested how FGF4 dose affected cell type proportions in ESCs. To be able to control FGF4 levels, we mutated the *Fgf4* gene in the GATA4-mCherry inducible cells. In contrast to the wild type control, *Fgf4* mutant cells continued to express high levels of NANOG, and showed almost no signs of differentiation after an 8 h doxycycline pulse followed by 40 h of culture in N2B27 medium alone (Figures 2C and 2D). Supplementing recombinant FGF4 after the doxycycline pulse rescued the differentiation of two discrete cell types that had similar NANOG- and GATA6-expression profiles as differentiating wild type cells (Figure S4). The proportions of these cell types depended on FGF4 concentration (Figures 2D and S4). Thus, FGF4 signaling and the cell-intrinsic transcriptional circuits underlying cell differentiation mutually regulate each other in a dose-dependent fashion, establishing communication via FGF4 as a potential mechanism for coordinating cell differentiation in the population.

We next sought to determine the spatial range of signaling via FGF4 ligands in ESCs. We first tested the role of global communication through FGF4 by comparing differentiation outcomes at different medium-to-cell ratios (Figure 2E). We reasoned that if FGF4 acted globally, ligand concentration would equilibrate in the medium, such that larger volumes would effectively reduce FGF4 concentration, and decrease the proportion of PrE-like in favor of Epi-like cells. In contrast to this expectation, cell type proportions changed negligibly with media volume, (Figure 2E), indicating that dilution of FGF4 ligands in the medium does not strongly affect cell type proportioning.

To test whether in contrast the communication is governed by local FGF4 signaling, we disrupted cell-cell contacts by trypsinizing and re-seeding cells at different densities immediately after doxycycline induction (Figure 2F). This treatment strongly reduced the proportion of PrE-like cells compared to the non-trypsinized control (compare second to three rightmost columns in Figure 2F). Furthermore, the proportion of PrE-like cells systematically increased with cell density (three rightmost columns in Figure 2F, p < 0.05). Together, these data suggest that cell-cell communication via FGF4 occurs locally and is positively influenced by cell-cell contacts.

To directly measure the spatial range of FGF4 signaling in ESC colonies, we seeded single wild type cells labelled with a constitutively expressed dsRed marker onto a layer of *Fgf4* mutant *Spry4*^*H2B-Venus*^ reporter cells (Morgani et al., 2018). 12 h after the addition of wild type cells, H2B-Venus was strongly expressed in a halo of reporter cells immediately surrounding the signal-emitting cells, but reporter expression dropped precipitously further away from them (Figure 2G). The spatial decay of the H2B-Venus signal was well approximated by a plateau of ∼ 14.4 µm (95% confidence interval 12.8 – 17.0 µm), followed by an exponential decay with a decay length of ∼ 11 µm (95% confidence interval 9.2 - 12.4 µm, Figure 2H). This is likely an overestimate of the immediate effective range of paracrine FGF4 signaling, as the transcriptional reporter integrates signaling activity over the entire duration of the experiment, during which cell divisions and movement will increase the distance between signal-sending and –receiving cells. Delaunay triangulation revealed that 16 h after the initiation of differentiation, the mean distance between a cell and its nearest and second-nearest neighbors was 14.0 ± 3.2 µm and 25.5 ± 5.3 µm, respectively (Methods, Figure S5). Thus, the range of cell-cell communication via FGF4 is spatially restricted and mainly couples nearest and second-nearest neighbors.

### Cell-cell communication via FGF4 underlies robustness of cell type proportions

We next asked if communication via FGF4 was the molecular mechanism underlying cell type proportioning. To this end, we titrated GATA4-mCherry induction levels, and compared the robustness of cell type proportions between wild type cells that can sense and secrete FGF4, and communication-deficient *Fgf4* mutant cells rescued with a fixed dose of recombinant FGF4 (Figures 3A - C). While cell type proportions remained relatively constant in the wild type (Figures 3 A - C, left column), they were strongly dependent on induction times - and hence initial GATA4-mCherry levels - in the rescued *Fgf4* mutant (right column). These broad deviations of cell type proportions in the mutant demonstrate that recursive communication via FGF4 underlies a population-level phenotype that is characterized by the differentiation of robust proportions of cell types from a wide range of starting conditions.

**Figure 3.**
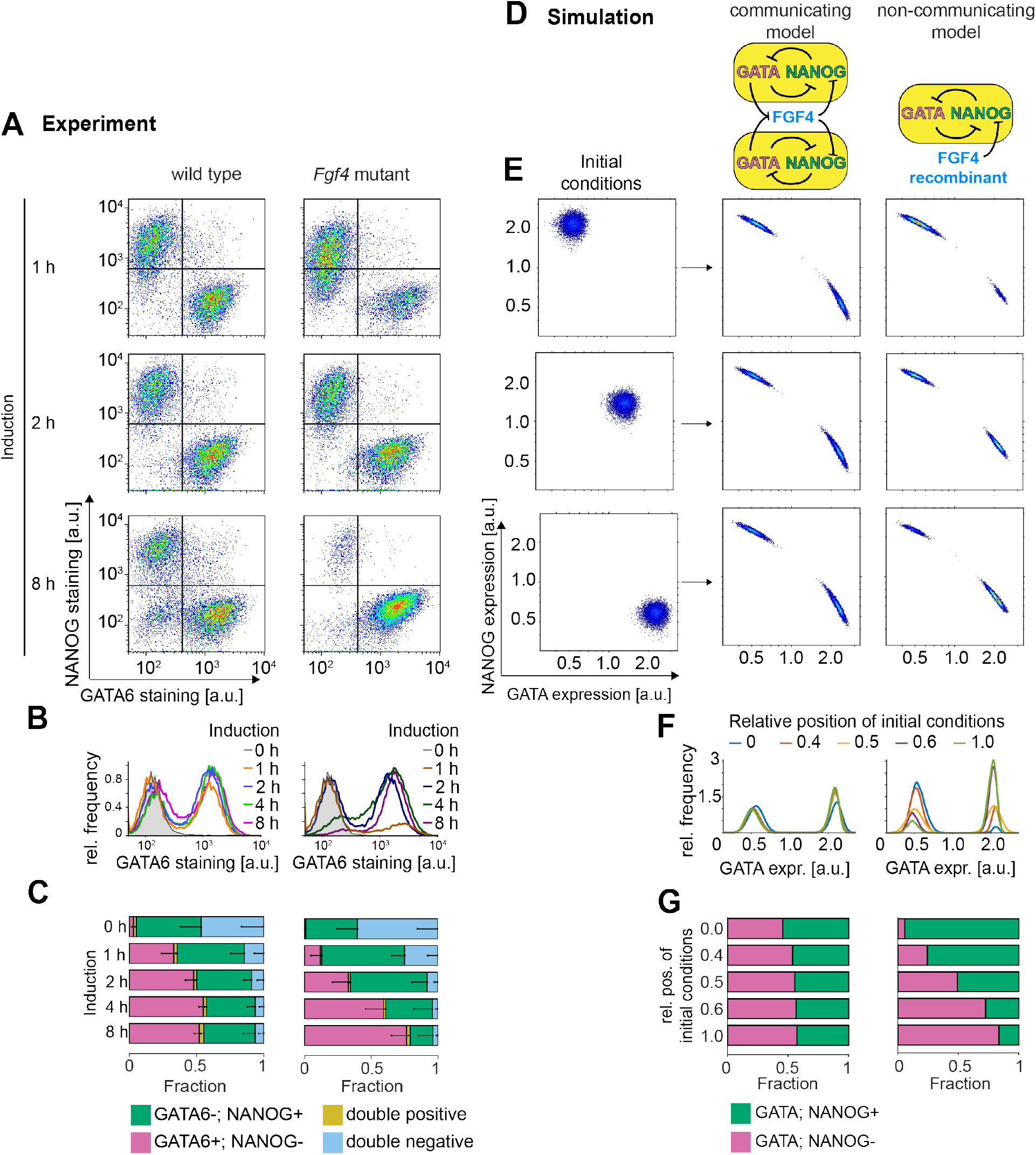
Cell-cell communication via FGF4 mediates cell type proportioning. **A – C** Cell differentiation in wild type (left) and Fgf4 mutant cells (right) after doxycycline pulses of variable length, followed by 40 h of differentiation in N2B27 alone (wild type) or in N2B27 supplemented with 10 ng/ml FGF4 (Fgf4 mutant). **A** Flow cytometry profiles depicting NANOG and GATA6 expression. Lines indicate gates to assign cell types. **B** Marginal distributions of GATA6 staining. **C** Quantification of average cell type proportions. GATA6+; NANOG-cells – magenta, GATA6-; NANOG+ cells - green, double positive cells - yellow, and double negative cells - blue. N = 4, error bars: 95% CI. **D** Schematic representation of the model with (left) and without cell-cell communication (right). **E** - **G** Influence of initial conditions (left column in **E**) on (GATA6+; NANOG-) and (GATA6-; NANOG+) cell type proportions in the model with (middle column) or without cell-cell communication (right column). **E** Distributions of initial conditions, shifting from NANOG+ to GATA+ (left column), and corresponding results of numerical simulations with (middle column) or without cell-cell communication (right column). **F** Respective GATA6 projections, equivalent to **B. G** Quantification of the cell type proportions obtained from the numerical simulations in **E**. See Figure S6 and Methods for model details and parameters.

To further explore how this population-level behavior of robust proportioning arises from cell-cell communication, we used numerical simulations to determine the dynamical properties of a communicating cell population. We developed a mathematical description by considering a previously characterized circuit consisting of mutually repressive interactions between GATA factors and NANOG (Schröter et al., 2015) that communicates among single cells through FGF signaling (Figure 3D, left). Specifically, we pose that GATA factors repress FGF4 expression (Figure 2), and that FGF signaling represses Nanog expression (Hamilton and Brickman, 2014; Schröter et al., 2015). Cell-cell communication was set between nearest and second-nearest neighbors in a population of N = 10000 cells on a 100 x 100 square grid (Figure S6A).

In the simulations, we considered several different scenarios of initial conditions, ranging from all cells being GATA-positive initially to all cells being NANOG-positive initially, to mimic the experimental settings (Figure 3E, left column). Within the coupled population, a stable proportion of two distinct gene expression patterns - (NANOG+, GATA-) or (NANOG-, GATA+) - was established (Figure 3E - G, middle column) from all starting conditions. In contrast, removing the communication in the model such that the dynamics of the mutually repressive NANOG-GATA circuit in single cells is only affected by a constant exogenous FGF input, the cell type proportions strongly depended on the initial conditions distributions (Figure 3D - G, right column). This congruence between the theoretical and experimental results indicates that recursive cell-cell communication via FGF signaling is sufficient to recapitulate our observation of robust cell type proportioning.

Bifurcation analysis of a minimal system of N = 2 communicating cells indicated that the robust proportioning of the two differentiated cell types in the wild type is a consequence of a new joint dynamical state of the system, characterized with an inhomogeneous steady state (IHSS, Figure S6B). An IHSS is generated at the population level, it is characterized by mutually exclusive gene expression patterns in the population, and has recently been proposed as a generic mechanism for robust cell type proportioning (Stanoev et al., 2019). This suggests that the emergence of an IHSS as a new dynamical solution in the population of FGF-coupled NANOG-GATA mutually repressive circuits is a possible dynamical basis for robust cell type proportioning in differentiating ESCs.

### Spatial organization of cell types indicates a shift from FGF4-dependent to FGF4-independent patterning mechanisms

The short spatial range of FGF4 signals in ESC cultures suggests that communication via FGF4 not only leads to robust global cell type proportions, but that the differentiated cell types should be also arranged in spatial patterns with local structure. We therefore analyzed the spatial arrangement of cell types at different time points after the initiation of differentiation. Staining for GATA6 and NANOG indicated that cell types were well mixed after 16 h and 24 h of differentiation, but clustering, particularly of GATA6-; NANOG+ cells, was observed after 40 h (Figure 4A). Analysis of the cell type composition of the immediate neighborhood of GATA6+; NANOG- (G+) and GATA6-; NANOG+ (N+) cells, identified with a 2D Gaussian mixture model (Figure S7, methods), corroborated a transition from well-mixed to clustered patterns (Figure S8A).

**Figure 4.**
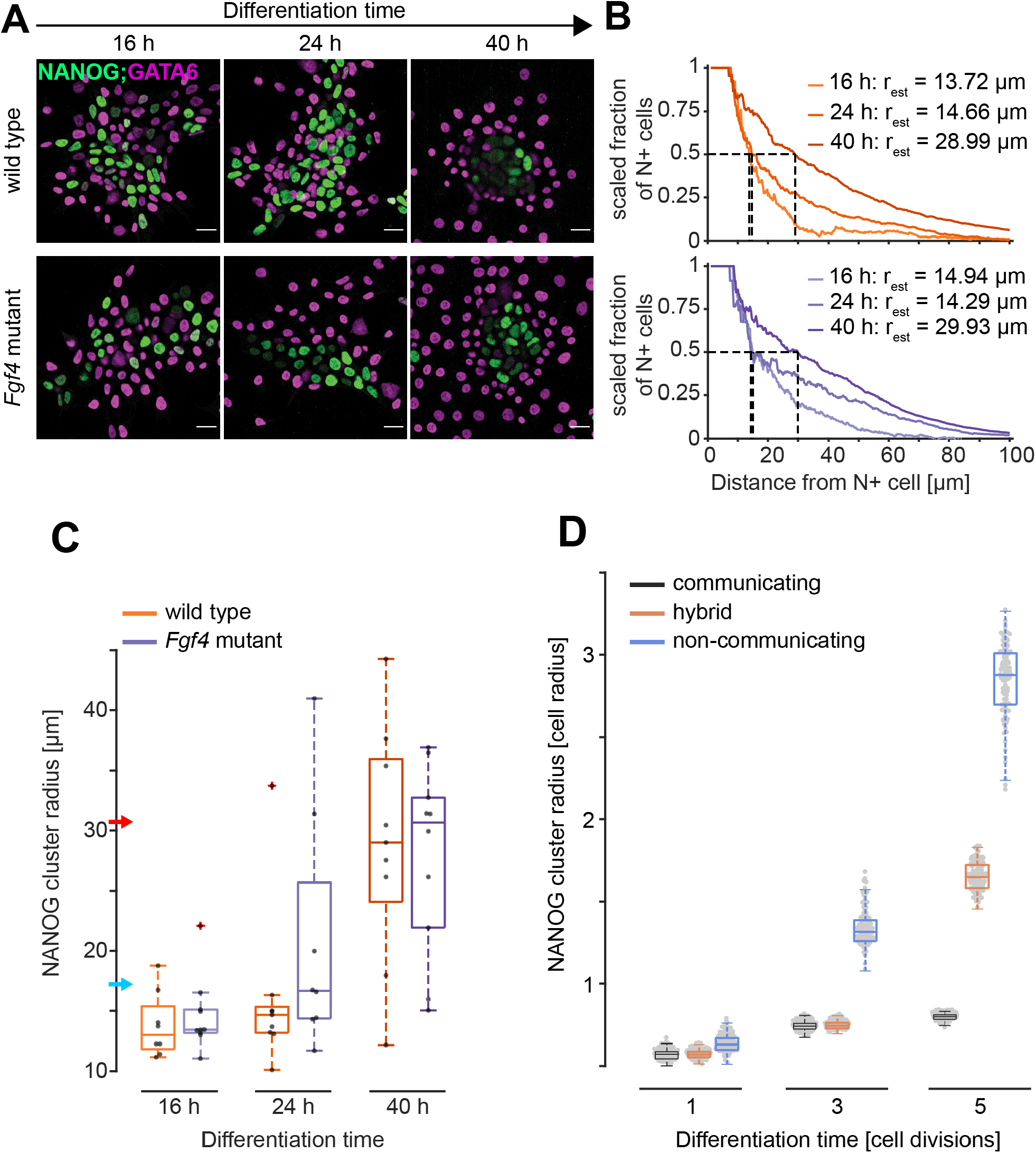
Spatial arrangement of cell types in wild type and Fgf4 mutant cells. **A** Immunostaining of wild type and Fgf4 mutant cells for NANOG (green) and GATA6 (magenta) at different time points after the initiation of differentiation. Wild type cells were induced with doxycycline for 8 h and differentiated in N2B27, Fgf4 mutant cells were induced for 4 h and differentiated in N2B27 supplemented with 10 ng/ml FGF4. **B** Estimation of NANOG cluster radius in a single field of view for wild type (orange, top) and Fgf4 mutant cells (blue, bottom) differentiated as in **A**. Graphs depict the scaled fraction of N+ cells within a specific radius around seed cells. Dashed lines indicate determination of cluster radius, the distance around NANOG+ cells from which the scaled fraction is equal to 0.5. **C** Summary statistics of quantified cluster radii for wild type (orange) and Fgf4 mutant cells (blue) differentiated as in **A**. Dots indicate values from individual fields of view (n ≥ 8), box plots show median, interquartile ranges and outliers (red cross). Blue and red arrowheads indicate mean distance between nearest and second nearest neighbors, respectively (Figure S5). **D** NANOG+ cluster radii quantified from 100 independent numerical realizations of the model with continuous communication (grey), without communication (blue), and of a hybrid model where communication is switched off after the third division (orange). Cluster radii were obtained after each cell cycle of a lineage tree simulation, before cell divisions take place.

Next, we set out to quantify cluster sizes during the differentiation time course. We first computed the scaled fraction of N+ cells in neighborhoods of increasing radius around all N+ cells in a field of view (Figure 4B, Methods). The value of the scaled fraction is 1 as long as all cells in the neighborhood are N+, and approaches zero when the composition of the local neighborhood equals the global composition of cell types. We defined the cluster radius as the distance around N+ cells at which this scaled fraction drops to 0.5 (dashed lines in Figure 4B).

The measure of cluster radius allows quantifying the length scales of spatially irregular features, however, its absolute value will be smaller than the physical size of the spatial features. In wild type cells, the median cluster radius was 13.0 µm and 14.7 µm at 16 h and 24 h of differentiation, respectively, and increased to 29.0 µm at 40 h of differentiation, corresponding to approximately 0.9, 1.0, and 2.1 cell diameters (Figure 4C, orange).

To investigate whether these experimentally determined wild type cluster patterns are consistent with short range signaling, we quantified the cell type spatial organization in model simulations. To be able to capture cluster formation due to cell division events, we considered a dividing cell population, where the grid size was doubled at each cell division event, and the daughter cells inherited gene expression states from the mother cell (Figure S9A). In the wild type case, the cluster radius increased only slightly, from 0.6 to 0.8 cell diameters over five divisions (Figures 4D, S9B, grey). This constant cluster size despite the state propagations during the cell divisions is maintained by the short-range communication that induces cell type transitions following each division in the simulation. The cluster radius in simulations is thus broadly consistent with the experimentally measured values at early, but not at late stages of differentiation.

Cluster formation at later stages of differentiation could be driven by long-range communication via FGF4, or by FGF4-independent mechanisms. To distinguish between these possibilities, we analyzed the spatial arrangement of cell types in rescued *Fgf4* mutant cells, using 4 h or 8 h of GATA4-mCherry induction together with differentiation in the presence of 10 ng/ml or 2.5 ng/ml FGF4, respectively, to obtain similar cell type proportions as in the wild type (Figures 2B and 3C). In both *Fgf4* mutant conditions, cell types were initially well mixed, and clustered at later stages of differentiation (Figures 4A and S8B – D). The median cluster radius in *Fgf4* mutants was similar to the wild type at 16 h, but then increased more rapidly until 24 h, to reach similar sizes as in the wild type at 40 h (Figures 4B, 4C, blue, and S10). The continuous increase in cluster size was recapitulated in simulations of the mutant case, where differentiation in the individual cells was initialized from a distribution of initial conditions. In these simulations, cluster sizes were initially comparable to the wild type case, but increased rapidly as cells divided, since in the absence of coupling-dependent cell type transitions gene expression states and thus cell types propagated locally (Figures 4D and S9, blue). Consequently, the transition from a well-mixed to a clustered cell type arrangement observed experimentally in the wild type could be recapitulated if following the settling to committed cell types, the communication was removed from the system (Figures 4D and S9, orange).

Taken together, these results suggest that at early stages, the observed spatial organization of cell types is consistent with regulation by a local cell-cell communication mechanism. Later on however, additional FGF4-independent mechanisms such as cell division and active cell sorting dominate the spatial organization.

### Heterogeneous differentiated cell types are maintained by intercellular communication

A central characteristic of a population-based mechanism for cell differentiation, such as the IHSS, is the interdependence of different cell types (Koseska and Bastiaens, 2017). It has been theoretically demonstrated that this property of the IHSS solution manifests in the regeneration of heterogeneous populations following separation of cell types after differentiation (Stanoev et al., 2019). We asked whether a similar behavior could be observed when PrE-like cells are isolated after both cell types have been established, using a live reporter for *Gata6* expression as a proxy to isolate PrE-like cells, and to subsequently monitor their differentiation state in different media (Figure 5A). Detection of both the up- and the downregulation of expression from the *Gata6* locus requires a short-lived transcriptional reporter protein. We therefore modified an established transcriptional reporter design (Freyer et al., 2015; Schröter et al., 2015) and knocked the coding sequence for the Venus-NLS-PEST protein (Abranches et al., 2013; Nagoshi et al., 2004) into the *Gata6* locus of a GATA4-mCherry inducible cell line.

**Figure 5.**
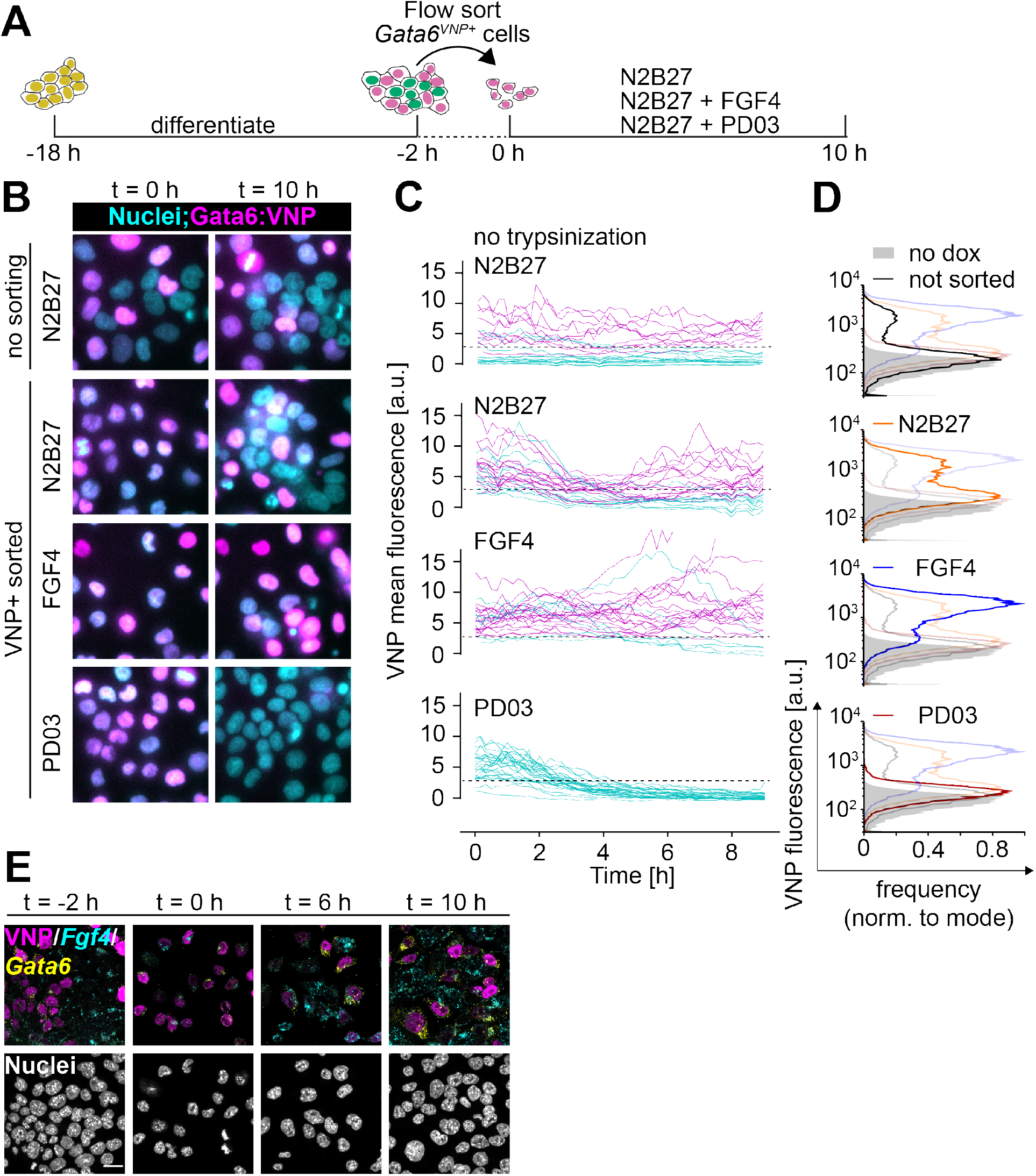
Heterogeneous cell identities are re-established by cell-cell communication. **A** Schematic representation of the cell-sorting experimental protocol. **B** Representative images of Gata6^VNP^ reporter expression in live cells of a non-trypsinized control (upper row) and in cells sorted for VNP expression. Left column is immediately after sorting, right column is after 10 h of culturing in N2B27 medium with the indicated supplements. **C** Live-cell traces of VNP expression in individual cells from non-trypsinized colonies (upper panel), or cells sorted for VNP expression upon culture in the indicated media. Traces are color coded according to expression levels at the end of the experiment. Dashed line indicates the threshold to separate putative VNP-positive cells (magenta) from VNP-negative cells (cyan). **D** Flow cytometry histograms of VNP expression of cells that had not been trypsinized and sorted (black, top), and of cells that had been sorted for VNP expression followed by 10 h of culture in the indicated media. Each panel shows the histogram of the relevant condition as a dark line, and distributions of all other conditions shaded for comparison. Non-induced control is in grey in all panels. **E** Staining for Fgf4 (cyan) and Gata6 (yellow) mRNA in Gata6^VNP^ reporter cells before sorting (left) and at 2 h, 6 h and 10 h after flow sorting of VNP-positive cells. VNP fluorescence in magenta. Scale bar, 20 µm.

Putative PrE-like cells were isolated by flow sorting for VNP expression 16 h after the end of a doxycycline pulse, a time-point when their spatial arrangement suggested that differentiation into heterogenous cell types is established by FGF4 communication. When cultured in N2B27 medium, sorted VNP-positive cells regenerated a mixture of VNP-positive and –negative cells within 10 h, resembling cell colonies that had not been disrupted and sorted (Figures 5B – D, first and second row).

Flow cytometry indicated a bimodal distribution of VNP expression both in unperturbed cultures and cultures that had been sorted for VNP-expression after 10 h of culture in N2B27 (Figure 5D). The smaller proportion of VNP-positive cells in the unperturbed cultures is likely due to insufficient induction of the GATA4-mCherry transgene in this cell line. The dynamics of VNP expression differed markedly between unperturbed cultures and sorted VNP-positive cells growing in N2B27: In unperturbed colonies, VNP expression was stable over time in most cells, while in sorted populations, the reporter was first globally downregulated before the heterogeneous expression patterns emerged (Figure 5C, first and second panel, Movies S1, S2). Similar transitions have been predicted *in silico* as a generic feature of the IHSS solution (Stanoev et al., 2019). The transient downregulation of VNP expression could be prevented by supplementing the culture medium with FGF4, which led to the maintenance of reporter expression in the majority of cells (Figures 5B – D, third row, Movie S3). Inhibition of FGF/ERK signaling with the MEK inhibitor PD03 in contrast resulted in the rapid downregulation of VNP expression following sorting in all cells (Figures 5B – D, fourth row, Movie S4). Reporter expression was likewise downregulated in sorted VNP-positive *Fgf4* mutant cells upon culture in N2B27 alone (Figure S11). These data indicate that cell-cell communication via FGF4/ERK regulates the re-establishment of a mixture of heterogeneous cell types in a population. The dynamics of *Fgf4* mRNA expression after sorting of VNP-positive cells and culture in N2B27 confirmed this (Figure 5E): Immediately after sorting, *Fgf4* transcripts could hardly be detected, as would be expected from the repression of *Fgf4* by endogenous GATA6 in VNP-positive cells. 6 h later, when the VNP reporter and hence endogenous GATA6 expression had dropped and repression was relieved, a subset of cells started to re-express *Fgf4* transcripts. 10 h after sorting, another subset of cells re-expressed *Gata6* mRNA and VNP, such that expression of *Fgf4* transcripts, *Gata6* mRNA and VNP in the population was mutually exclusive, similar to the situation before sorting. Taken together, these results indicate FGF/ERK signaling re-establishes populations with different cell types following the isolation of PrE-like cells. In unperturbed cell colonies, intercellular communication via FGF/ERK therefore not only generates, but also actively maintains balanced proportions of differentiated cell types.

## Discussion

Here we report emergent population-level behavior during the differentiation of Epi- and PrE-like cells from ESCs expressing inducible GATA factors: Robust proportions of the two cell types are specified from a wide range of GATA induction levels, and re-established from isolated PrE-like cells. This emergent global behavior relies on local cell-cell communication via FGF4. The observed differentiation characteristics recapitulate the properties of a population-based dynamical solution, an inhomogenous steady state, recently proposed as a generic mechanism underlying robust differentiation (Stanoev et al., 2019). Our results suggest a new function for FGF signaling, which is to generate and maintain robust proportions of differentiated cell types.

The specification of Epi- and PrE-like cells in ESCs shows both molecular and functional parallels to the patterning of the ICM of the mouse preimplantation embryo. In contrast to previous studies that reported PrE-like differentiation in *Fgf4* mutant ESCs upon permanent high-level expression of exogenous GATA factors (Kang et al., 2012; Wamaitha et al., 2015), we find that PrE-like cells do not differentiate from *Fgf4* mutant ESCs upon transient GATA induction. This recapitulates the *Fgf4* mutant phenotype in the embryo (Feldman et al., 1995; Kang et al., 2012; Krawchuk et al., 2013). In both *Fgf4* mutant ESCs and embryos, cell type proportions can be controlled by recombinant FGF4 in a dose-dependent manner (Yamanaka et al., 2010; Krawchuk et al., 2013). Furthermore, the differentiation of Epi- and PrE-like cells *in vitro* recapitulates the remarkably constant proportions of cell types seen in the developing embryo (Saiz et al., 2016). Lastly, cell identities in ESC populations are plastic and can be re-specified upon changing a cell’s environment, again similar to observations in the embryo (Grabarek et al., 2012; Martinez Arias et al., 2013). A recent study using chimaeras and targeted ablation of specific cell types concluded that an FGF4-based population-level mechanism balances the size of the Epi and the PrE lineage in the mouse embryo (Saiz et al., 2020). We used doxycycline-inducible transgenes in wild type and *Fgf4* mutant ESCs as an orthogonal experimental system to demonstrate that a similar molecular and dynamical mechanism operates during cell differentiation *in vitro*. In addition to the previous observations, the ESC system also allowed us to show that cell type proportions can be robustly established from a broad range of initial conditions. The first direct measurements of the spatial range of cell-cell communication via FGF4 in differentiating ESCs finally provides indications how the spatial arrangement of cell types is established during development.

The short signaling range of FGF4 during the differentiation of Epi- and PrE-like cells *in vitro*, together with the progressive FGF-independent spatial clustering once cell types have been specified, suggest an evolutionary explanation for the spatially random differentiation of Epi and PrE cells in the ICM (Chazaud et al., 2006; Plusa et al., 2008; Rossant et al., 2003). Rather than settling on a mechanism that produces the end result of spatially separated tissues in one step, evolution might have taken a “tinkering” approach (Jacob, 1977), combining communication via FGF4 as a mechanism that allows specifying robust proportions of cell types, with a subsequent sorting step, a sequence that is conserved *in vitro*.

FGF4 signaling during the differentiation of Epi- and PrE-like cells shows several parallels to general properties of the Delta-Notch signaling system. The signaling range of FGF4 is spatially restricted, akin to the juxtacrine signaling mode of Delta-Notch. Communication via FGF4 in differentiating ESCs establishes and maintains discrete cell types through repressive coupling-signals sent by one cell type repress the identity of that same type in neighboring cells – much like Delta-Notch signaling during lateral inhibition (Ferrell, 2012; Henrique and Schweisguth, 2019; Hori et al., 2013; Simpson, 1990). The short spatial range of the FGF4 signal requires cell types to be well mixed immediately after specification, similar to the spatial patterning of cell types arising from Delta-Notch-mediated lateral inhibition (Collier et al., 1996). Consequently, several hallmarks of the emergent population-level behavior established by FGF4 in differentiating ESCs have been observed in an engineered cell system where cells communicate via Delta-Notch, such as the differentiation of discrete cell types in reproducible proportions, the re-establishment of those proportions upon removal of one cell type, and the dependence of cell type proportions on cell density or contact (Matsuda et al., 2015). Thus, when embedded in appropriate intracellular regulatory circuits, molecularly diverse intercellular communication systems can yield similar functional outputs.

In rescued *Fgf4* mutant cells that do not communicate, differentiation outcomes in the cell population strongly depend on the distribution of GATA4-mCherry induction levels. This is in line with predictions from single cell models for cell differentiation, in which initial conditions in individual cells strongly influence their differentiation path (Huang et al., 2007). When cells communicate via FGF4 in contrast, the collective differentiation outcome is robust and becomes independent from the distribution of GATA4-mCherry induction levels. Thus, the behavior of the communicating cell population cannot directly be extrapolated from the behavior of single isolated cells.

Theoretically, the conceptual differences between single-cell and population-based modes of differentiation manifest in the emergence of a new type of solution that jointly describes the heterogeneous cell identities, an IHSS, in the communicating cell population. The robust generation of cell type proportions irrespective of initial conditions, and their active maintenance through intercellular communication that we observe experimentally, are two key properties of the IHSS (Stanoev et al., 2019). This suggests that the IHSS is a likely dynamical mechanism underlying the differentiation of cells with discrete identities during mammalian preimplantation development. Given the pervasiveness of robust cell type proportioning during development and homeostasis (Viader-Llargués et al., 2018), it is likely that similar population-based mechanisms underlie canalized development in diverse systems in which multipotent progenitor cells give rise to several differentiated cell types.

## Methods

### Cell lines

Cell lines used in this study were E14tg2a (Hooper et al., 1987) and an *Fgf4* mutant *Spry4*^*H2B- Venus/+*^ line that we have previously described (Morgani et al., 2018). dsRed-labelled cells were from an E14tg2a-background and kindly supplied by J Nichols. The Gata6:VNP reporter was established in the background of a line carrying a doxycycline-inducible GATA4-mCherry transgene in the Col1a1 locus as well as a randomly integrated H2B-Cerulean nuclear marker driven by a CAGS promoter described in (Schröter et al., 2015).

E14tg2a-based inducible cell lines were maintained on fibronectin-coated tissue culture plastic in 2i + LIF medium, which consists of a N2B27 basal medium supplemented with 3 µM CHIR99021 (Tocris), 1µM PD0325901 (SelleckChem) and 10 ng/ml LIF (protein expression facility, MPI Dortmund). For maintenance of *Fgf4* mutant subclones, we supplemented the 2i + LIF medium with 10% fetal bovine serum (FBS), as *Fgf4* mutant lines showed severely decreased proliferation upon long-term culture in 2i + LIF alone. FBS was removed at least one day before the experiment.

*Spry4*-reporter cell lines to measure signaling range, as well as *Gata6*-reporter cell lines were maintained on gelatin coated dishes in GMEM-based medium supplemented with 10% FBS, sodium pyruvate, 50 µM β-mercaptoethanol, glutamax, non-essential amino acids and 10 ng/ml LIF. 1 µM PD0325901 was added to the cultures of *Spry4*- and *Gata6*-reporters three days before the experiment, to downregulate *Spry4* reporter expression, or to capacitate cells for PrE-like differentiation (Schröter et al., 2015).

FGF4 was from Peprotech and supplied in the indicated concentrations, together with 1 µg/ml heparin (Sigma).

### Genetic engineering of ESC lines

Doxycycline-inducible *GATA4*-mCherry inducible ES cells were generated by electroporation of 50.000 E14tg2a ES cells with 4 µg of pPB-TET-GATA4-mCherry, 4 µg pCAG-rtTA-Neo, and 4 µg pCAG-PBase (Wang et al., 2008) followed by G418 selection (400 µg/ml) one day after transfection. We established more than 10 independent clonal lines and assayed induction levels and homogeneity by flow cytometry 2 – 8 h after induction of transgene expression by adding 500 ng/ml doxycycline to the culture medium. Four clones with homogeneous induction levels were chosen and maintained under G418 selection, to circumvent silencing of the inducible transgene.

Mutagenesis of the *Fgf4* locus was performed as previously described (Morgani et al., 2018). *Fgf4* loss of function clones were identified by PCR-amplification, cloning and sequencing of a sequence around the *Fgf4* start codon. We either selected clones with a targeted mutation delivered by a single-stranded DNA repair template that we have previously shown to disrupt *Fgf4* function (Morgani et al., 2018), or selected at least two independent clones carrying indels around the start codon that introduced frameshift as well as nonsense mutations. All independent clones with random indels showed indistinguishable behavior in the differentiation assays.

The *Gata6* reporter cell line was generated using previously described knock-out first targeting arms of the EUCOMM project (Skarnes et al., 2011), combined with a VNP reporter cassette (Nagoshi et al., 2004) and a neomycin resistance gene driven from a human b-actin promoter. This construct was integrated by homologous recombination into a line carrying a doxycycline-inducible GATA4-mCherry transgene in the *Col1a1* locus as well as a randomly integrated H2B-Cerulean nuclear marker driven by a CAGS promoter described in (Schröter et al., 2015). Clones were screened for correct integration of the reporter construct by long range PCR spanning the targeting arms.

The targeting construct to generate the *Spry4*^*H2B-Venus*^ allele in GATA4-mCherry inducible cell lines was based on the one used in (Morgani et al., 2018), except that the puromycin selectable marker was exchanged for a neomycin cassette. The construct was integrated into ESCs by homologous recombination. To increase targeting efficiency, cells were co-transfected with a plasmid expressing Cas9 and a sgRNA that targets a sequence at the 5’ end of the 5’ targeting arm which is present in the endogenous *Spry4* locus, but not the targeting construct. Neomycin-resistant clones were expanded and screened for correct integration of the reporter construct by long range PCR spanning the targeting arms. All genetically modified lines were karyotyped using standard procedures (Nagy et al., 2008), and only lines with a median chromosome count of n = 40 were used for experiments.

### Immunostaining and image analysis

Immunostaining of adherent cells was performed as previously described (Schröter et al., 2015). Antibodies used were anti-NANOG (e-bioscience, eBioMLC-51, 14-5761-80, final concentration 2.5 µg/ml), anti-GATA6 (R&D AF1700, final concentration 1 µg/ml), and anti-FLAG (Sigma-Aldrich F1804-200, final concentration 1 µg/ml). Secondary antibodies were from Invitrogen/LifeTech. Images were acquired using a 63x 1.4 N.A. oil-immersion objective on a confocal Leica SP8 microscope, with all settings held constant between replicates. Images were quantified using custom scripts written for ImageJ (NIH) and in Matlab (The Mathworks).

### In situ HCR

Probe sets for *Nanog, Gata6* and *Fgf4* and corresponding Alexafluor-labelled amplifiers for staining of mRNA molecules via third generation in situ HCR (Choi et al., 2018) were sourced from Molecular Instruments. Staining was performed according to manufacturer’s instructions. Briefly, adherent cells were fixed for 15 minutes with 4% paraformaldehyde, washed with PBS and permeabilized for several hours in 70% ethanol at −20°C. Cells were then washed twice with 2x SSC and equilibrated in probe hybridization buffer for at least 30 minutes. Transcript-specific probes were used at a concentration of 4 nM and hybridized overnight. Excess probe was removed through several washes with probe wash buffer and 5x SSCT, and cells were equilibrated in amplification buffer for at least 30 minutes. Fluorescently labeled amplifiers were used at a concentration of 60 nM. Amplification was allowed to proceed for 16 – 24 hours at room temperature. Excess amplifier was removed by several washes with 5x SSCT, followed by counterstaining with Hoechst 33342 and mounting in glycerol-based medium. Imaging was performed on an SP8 confocal microscope with a 63x (NA1.4) lens.

### Flow cytometry

Staining for flow cytometric analysis of intracellular antigens was performed as previously described (Schröter et al., 2015). Primary and secondary antibodies were the same as used for immunostaining. mCherry fluorescence measurements and cell sorting were performed on a BD FACS Aria. All other flow cytometric analysis was carried out using a BD LSR II. Single cell events were gated based on forward and side scatter properties. GATA4-mCherry expression measurements were normalized to the respective uninduced control. Gates to separate marker-positive from marker-negative cells were determined visually as the threshold that best bisected the bimodal distribution of marker expression across all samples within one experiment.

### Decay length measurements

*Fgf4* mutant Spry4^H2B-Venus^ reporter cells (Morgani et al., 2018) were seeded at a density of 5∗10^4^ cells/cm^2^ in N2B27. 2 hours later, dsRed-expressing cells were added at a density of 500 cells/cm^2^. For the first 3 hours of co-culture, the medium was supplemented with 250 to 500 nM siR-Hoechst (Lukinavičius et al., 2015) to label nuclei. 12 h later, live cells were imaged on a Leica SP8 confocal system. Nuclei were segmented in FIJI (Schindelin et al., 2012), and for each Spry4^H2B-Venus^ reporter cell in the vicinity of a ds-Red expressing cell, the background-subtracted Venus fluorescence intensity as well as the distance to the center of mass of the dsRed expressing cells was determined. Cells were grouped according to their distance from dsRed expressing cells in 3 µm bins, and mean fluorescence intensities for each bin plotted versus their distance. Decay length was estimated in GraphPad Prism by fitting a plateau followed by a one-phase decay function.

### Determination of cell-cell distances

Neighborhood graphs were constructed for each field of view by using the cell positions to generate a Delauney triangulation and the corresponding Voronoi diagram. Spurious links between non-adjacent cells were trimmed by excluding links in the Delauney graph that do not directly pass through the shared Voronoi edge between the two respective Voronoi cells. Links between adjacent cells (purple links in Fig S5A) were pooled to generate a distance distribution between nearest neighbors (purple histogram, Fig S5B). The distances of links between unconnected cells that share a neighbor (yellow in Fig S5A), were likewise pooled to give the distribution of second-nearest neighbor distances (yellow histogram, Fig S5B). Both distributions were independently fit with a 2-component Gaussian mixture model, to separate the true distributions of nearest and second-nearest neighbor distances from higher-order distributions arising from erroneously assigned links (purple and yellow dashed lines, Fig S5B). The mean and SD of nearest- and second-nearest neighbor distances was estimated from the first component of these Gaussian fits.

### Assignment of cell types with Gaussian mixture model

For spatial clustering analysis, we focused on GATA6+; NANOG-(G+) and GATA6-; NANOG+ (N+) cells only. To identify these cell types specifically, we applied a two-component Gaussian mixture model (GMM) fit to single cell distributions in the GATA6; NANOG expression space, using the MATLAB function fitgmdist(). The fitted GMM assigns each cell a posterior probability of its association with one of the two component distributions. Cells with a posterior probability of >0.9 for one of the two components were classified to the corresponding cell type.

### Analysis of neighborhood composition

To determine local neighborhood composition, we first generated a matrix of Euclidean distances between every cell classified as either G+ or N+ for every individual field of view. For every cell, we then computed the fraction of G+ and N+ cells within a Euclidean distance of 31.7 µm, encompassing most of the nearest and second nearest neighbors. Random distributions (blue lines in Figure S8) were calculated by populating the local neighborhood of each cell randomly from the global distribution in each treatment.

### Measurement of cluster radius

The average cluster radius of N+ cells was estimated with the same method in experimental data where cell types had been categorized with a Gaussian mixture model, and in simulations of the model. We calculated for all N+ cells in a field of view or simulation run the decrease in the fraction of N+ cells in neighborhoods of increasing radius, settling to the overall fraction of N+ cells. For scaling, we first subtracted the overall fraction of N+ cells in the respective field of view or simulation run, and then normalized values such that the scaled fraction at zero distance was set to one again. The cluster radius is the distance at which the scaled fraction of N+ cells is equal to 0.5.

### Statistical analysis

GraphPad Prism was used to perform significance testing (ANOVA).

### Live cell imaging and tracking

To track *Gata6* reporter expression in live cells, PrE-like differentiation was induced by a 6 h pulse of doxycycline-treatment in serum-containing medium as described in (Schröter et al., 2015). 16 hours after doxycycline-removal, cells were either switched directly to N2B27 medium lacking phenol red, or trypsinized, sorted for reporter expression, and seeded on fibronectin-coated imaging dishes (ibidi µ-slides). Time-lapse imaging was started within 2 h after sorting on an Olympus IX81 widefield microscope equipped with LED illumination (pE4000, CoolLED) and a Hamamatsu c9100-13 EMCCD camera. Hardware was controlled by MicroManager software (Edelstein et al., 2001). Time-lapse movies were acquired using a 40x oil immersion lens (NA 1.2), with 10-minute time intervals.

Cell tracking was carried out with TrackMate (Tinevez et al., 2017) based on the constitutively expressed H2B-Cerulean nuclear marker. Fluorescence intensity was measured in a circular region of interest in the center of the nucleus, and background-subtracted fluorescence intensities plotted in Python. Trace color in Figure 4D was assigned according to fluorescence intensity in the last frame of the movie, with respect to the estimated intensity threshold used for flow sorting (dashed line).

### Computational model for cell type proportioning

The model of the intercellular communication system (Figure 3D) is adapted from Stanoev et al. (2019), and is described with the following set of equations: 

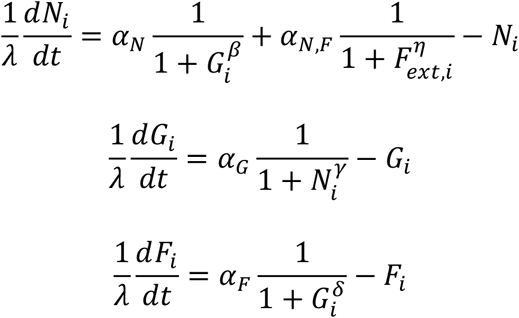

*N*_*i*_ and *G*_*i*_ describe NANOG and GATA6 protein expression levels in cell *i*, regulated by mutual inhibition, while *F*_*i*_ is the secreted FGF4 whose production is downregulated by GATA6. 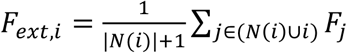 is the extracellular FGF4 concentration that is sensed by cell *i* from its neighborhood *N*(*i*), resulting in downregulation of NANOG production in the cell. *α*_*N*_ = 2.5, *α*_*N,F*_ = 0.5, *α*_*G*_ = 3 and *α*_*F*_ = 3 denote production rate constants, *β* = *η* = *γ* = *δ* = 2 are the Hill coefficients, degradation rates were set to 1 as *λ* = 50 was used as a scaling kinetic parameter. 10000 cells were deployed on a regular 100×100 two-dimensional lattice with no-flux boundary conditions. Cell-cell communication was modeled to be short-range, reflecting the experimental wild-type case, i.e. communication between direct neighbors and cells two hops away on the lattice (Figure S6A). When mimicking the *Fgf4* mutant case, communication between cells was excluded, and an external input was modeled with *F*_*ext*_ = 1.2.

The cell populations were initiated analogously to the experimental case, by varying the initial conditions of all cells from being NANOG-expressing, through intermediate NANOG and GATA6 expression, to being GATA6-expressing (Figure 3E, left column). More specifically, the variables were sampled independently from unimodal Gaussian distributions *𝒩* _*ics*_ (*p*), *σ*_*ics*_ = 0.1 ∗ *μ*_*ics*_ (*p*), with the mean *μ*_*ics*_ (*p*) = (1 – *p*) ∗ *μ*_*G–;N+*_ + *p* ∗ *μ*_*G+;N–*_ placed on the line segment connecting the GATA6-; NANOG+ state *μ*_*G–;N+*_ and the GATA6+; NANOG-state *μ*_*G–;N+*_, partitioning it in proportion *p. p* ∈ {0, 0.4, 0.5, 0.6, 1} was used for the quantifications in Figure 3G. Samples from around the endpoints and the midpoint 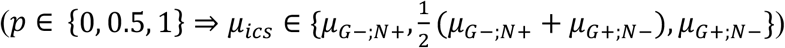are shown in Figure 3E, left column.

Cell heterogeneity was introduced by varying all of the parameters independently with standard deviation of 0.02 from the respective values for each cell. Stochastic differential equation model was constructed from the deterministic equations by adding a multiplicative noise term *σXdW*_2_, where *dW*_*t*_ is the Brownian motion term, *X* is the variable state and *σ* = 0.1 is the noise term. The model was solved with Δ*t* = 0.01 using the Milshtein method (Milshtein, 1974). Following integration, cell identities were estimated by comparing the NANOG and GATA6 values from the final states of the cells, and the ratios were computed.

For comparing the spatial organizations between communicating and non-communicating cells at different time points, periodic synchronous cell divisions in the population were included in the model as in (Stanoev et al., 2019), spanning 5 cell cycles. Cell divisions occur along the horizontal and vertical axes on the grid alternately, sequentially yielding lattices of 10×10, 10×20, 20×20, 20×40 and 40×40 (Figure S9A). At every cell division, the final state of the mother cell is passed on to the daughter cells’ initial conditions. The mother cell’s parameter set is also inherited. Spatial organizations were analyzed at the end of each cell cycle, after the collective state is allowed to reach a steady state in a deterministic fashion, by estimating the N+ cluster radius as described above.

For the hybrid model, it was assumed that after the third cell cycle the cells commit to their current fates and the communication becomes inconsequential, effectively bringing about a switch to a non-communicating grid. For all conditions, cells’ states were initialized with Gaussians with *μ*_*ics*_ (0.5), as described above.

## Acknowledgements

We thank J. Nichols for sharing dsRed-labelled ESCs, L. Süther for help with generating inducible cell lines, M. Schulz and S. Müller for assistance with microscopy and flow cytometry, and P. Bastiaens for stimulating discussion and conceptual input on the project. We thank P. Bieling, J. de Navascues, G. Vader, S. Fischer and the members of the Schröter and Koseska groups for discussions and comments on the manuscript. All authors are supported by the Max Planck Society.

## Author contributions

Conceptualization, A.K. and C.S.; Methodology, A.S. and C.S.; Investigation, D.R., A.B., A.S., M.P., and C.S.; Validation, D.R., A.B., and M.P.; Visualization, D.R. and A.B.; Formal analysis, D.R. and A.B.; Writing – original draft, C.S.; Writing – review and editing, all authors; Supervision, A.K. and C.S..

## Supplementary Figures

**Figure S1.**
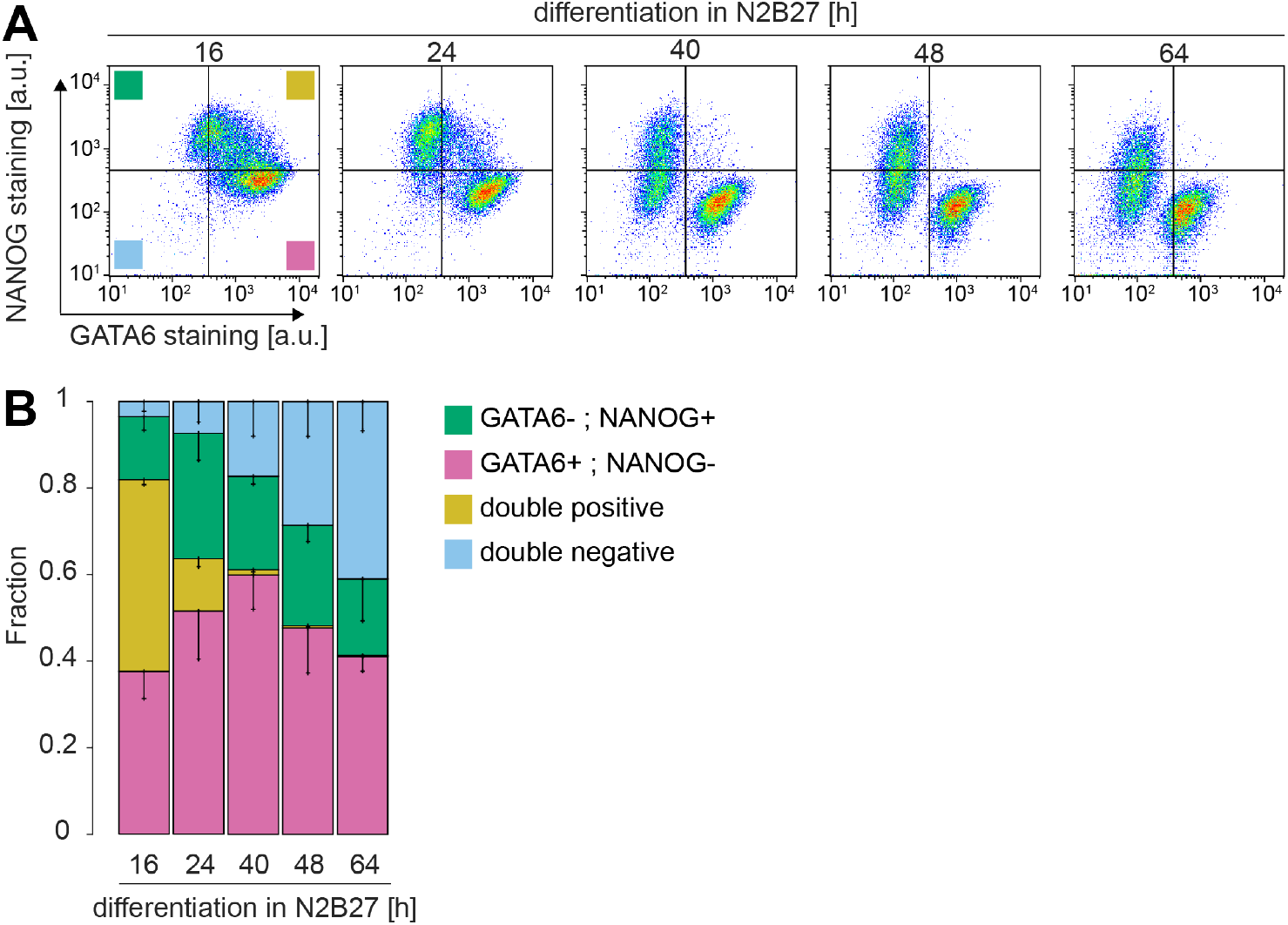
Time course of cell type proportions following pulsed GATA4-mCherry expression. **A** Flow cytometry profiles of cells stained for NANOG and GATA6 after an 8 h doxycycline pulse to induce GATA4-mCherry expression, followed by differentiation in N2B27 medium for the indicated durations. Two clusters corresponding to NANOG-high; GATA6-low Epi-like cells and NANOG-low, GATA6-high PrE-like cells can already be distinguished after 16 h of differentiation, but are only fully separated at 40 h and beyond. Solid lines and color code in leftmost panel indicate gating strategy to assign cell types. **B** Average cell type proportions from N = 3 independent experiments. Fraction of GATA6+, NANOG-cells in magenta, GATA6-, NANOG+ cells in green, double positive cells in yellow, and double negative cells in blue. Error bars: 95% CI.

**Figure S2.**
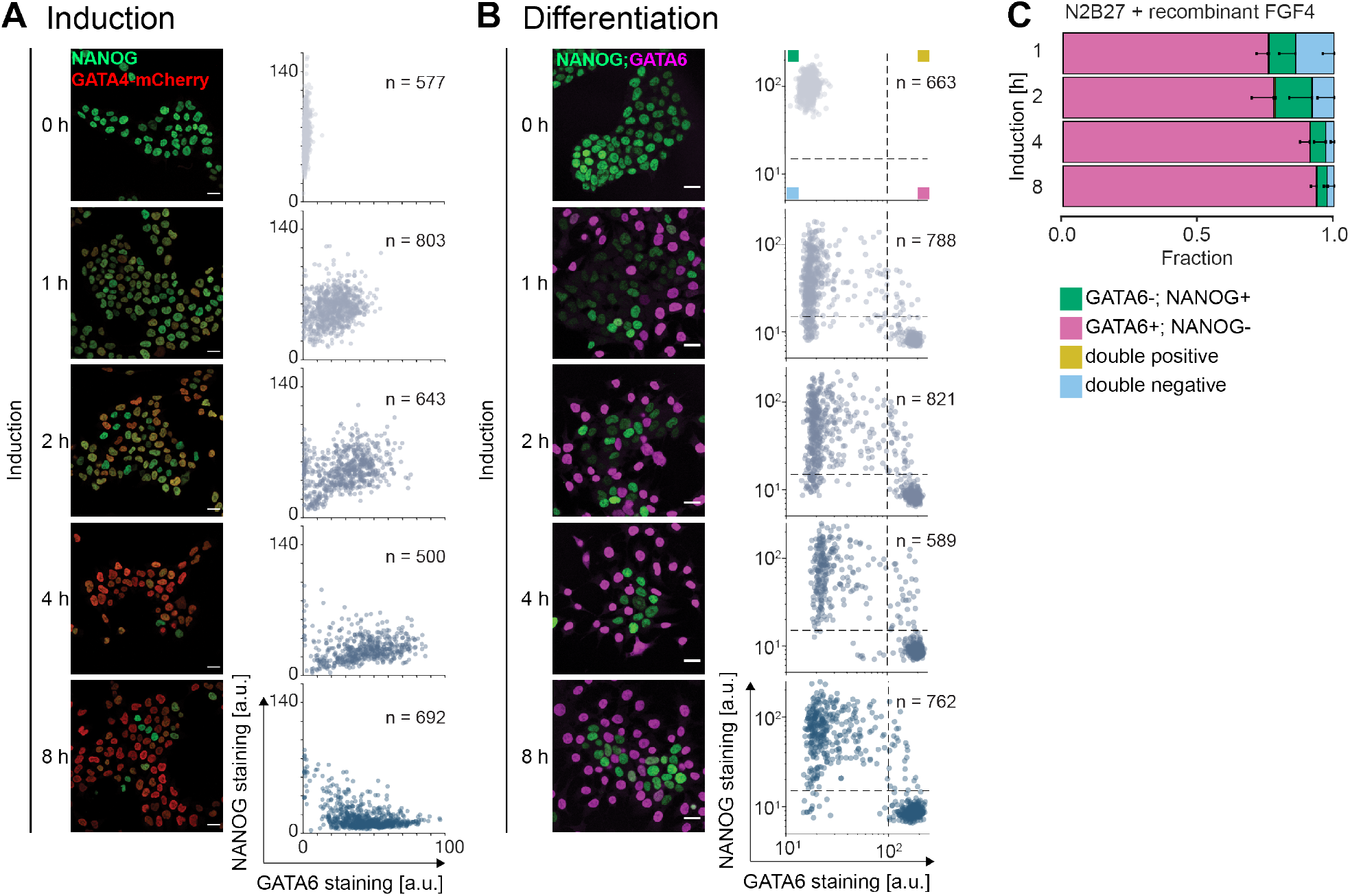
Quantitative immunofluorescence analysis of NANOG and GATA expression. **A** Immunostaining (left) and single-cell quantification (right) of NANOG (green) and GATA4-mCherry (red) expression in inducible cells after indicated durations of doxycycline treatment in 2i + LIF medium. **B** Immunostaining (left) and single-cell quantification (right) of NANOG (green) and GATA6 (magenta) expression in inducible cells after indicated durations of doxycycline treatment, followed by 40 h differentiation in N2B27 medium. Immunofluorescence micrographs in **A** and **B** for 0, 2, and 8 h of doxycycline induction are reproduced from Fig. 1 for comparison. Dashed lines in scatter plots in **B**: Thresholds to assign cell types; upper left quadrant: GATA6-; NANOG+; lower right quadrant GATA6+; NANOG-; upper right quadrant double positive; lower left quadrant double negative. Scale bars, 20 µm. **C** Proportions of cell types upon indicated durations of doxycycline induction followed by 40 h of differentiation in N2B27 medium supplemented with 10 ng/ml FGF4. Cell identities were determined by immunostaining and quantitative immunofluorescence. Fraction of GATA6+, NANOG-cells in magenta, GATA6-, NANOG+ cells in green, double positive cells in yellow, and double negative cells in blue. N = 4, error bars: 95% CI.

**Figure S3.**
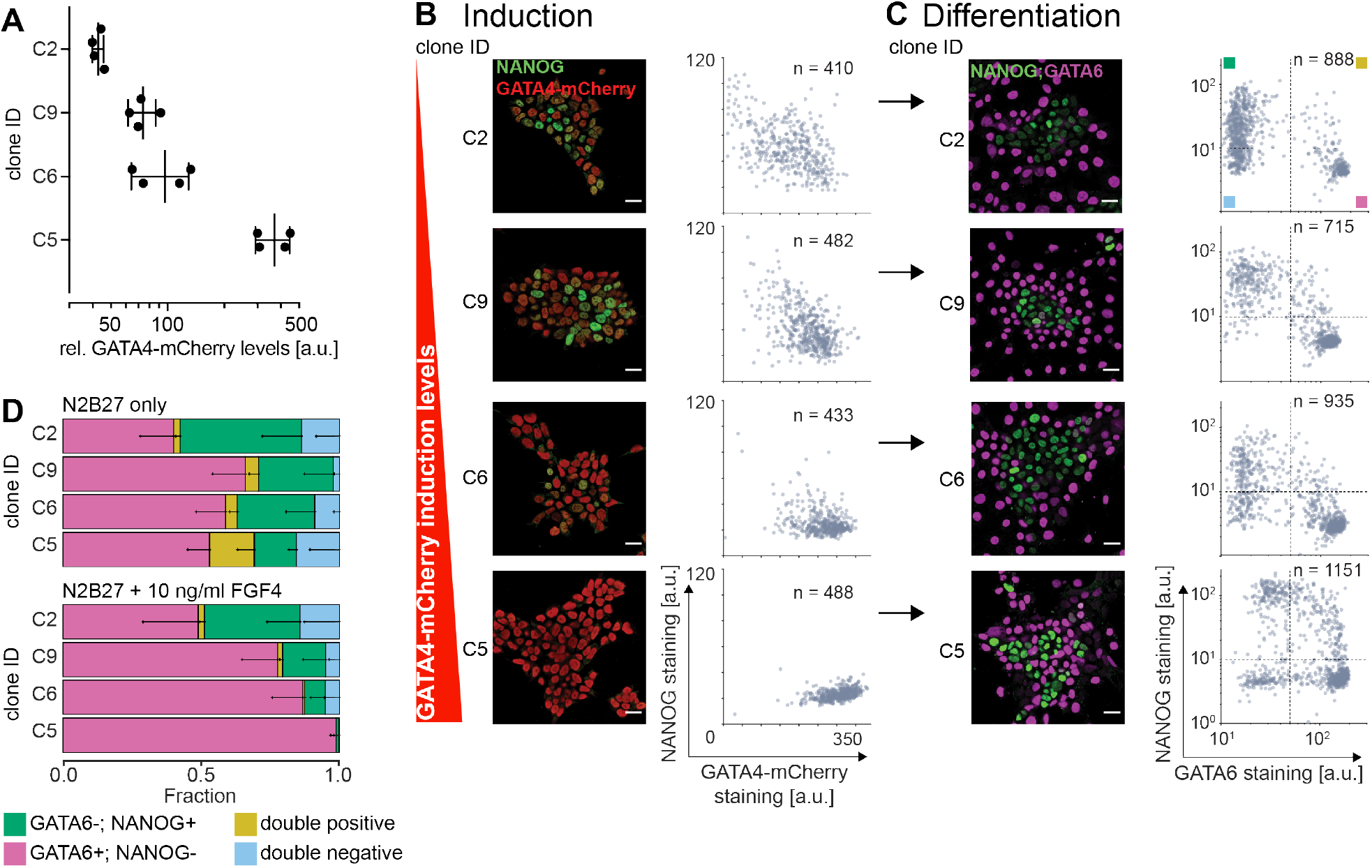
Robust cell type proportioning in independent clonal cell lines. **A** GATA4-mCherry induction levels in four independent clonal inducible cell lines after 8 h of doxycycline treatment in 2i + LIF medium, measured by flow cytometry. Clones are ordered by GATA4-mCherry expression strength, fluorescence values were normalized to non-induced control cells. Plot shows individual data points and mean ± SD from N = 4 independent experiments. **B** Immunostaining (left) and single-cell quantification(right) of NANOG (green) and GATA4-mCherry (red) expression in the same clonal lines analyzed in **A** after 8 h of doxycycline stimulation in 2i + LIF medium. **C** Immunostaining (left) and single cell quantification (right) of NANOG (green) and GATA6 (magenta) expression in cells from independent clonal lines treated with doxycycline for indicated periods of time and differentiated in N2B27 for 40 h. Dashed lines: Thresholds to determine cell types; upper left quadrant: GATA6-; NANOG+; lower right quadrant GATA6+; NANOG-; upper right quadrant double positive; lower left quadrant double negative. Clones are ordered by GATA4-mCherry induction strength as in **A**. Scale bars in **A, C**, 20 µm. **D** Average cell type proportions in clonal lines differentiated as in **C** in N2B27 only (top) or in N2B27 supplemented with 10 ng/ml FGF4 (bottom).Fraction of GATA6+, NANOG-cells in magenta, GATA6-, NANOG+ cells in green, double positive cells in yellow, and double negative cells in blue. N = 4, error bars: 95% CI.

**Figure S4.**
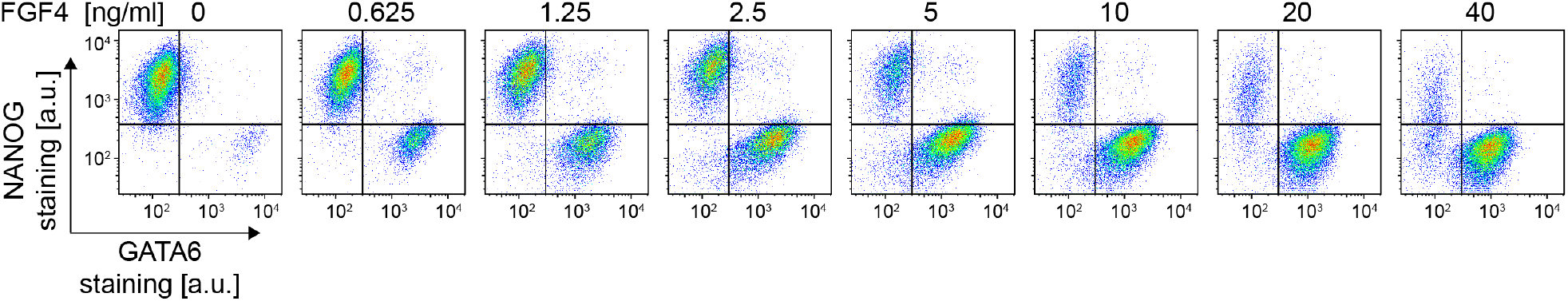
FGF4 dose regulates cell type proportions in Fgf4 mutant cells. Flow cytometry profiles of Fgf4 mutant cells stained for NANOG and GATA6 after 8 h of doxycycline induction, followed by 40 h of differentiation in N2B27 medium supplemented with 1 µg/ml heparin and the indicated concentrations of recombinant FGF4. Lines indicate gates to assign cell types.

**Figure S5.**
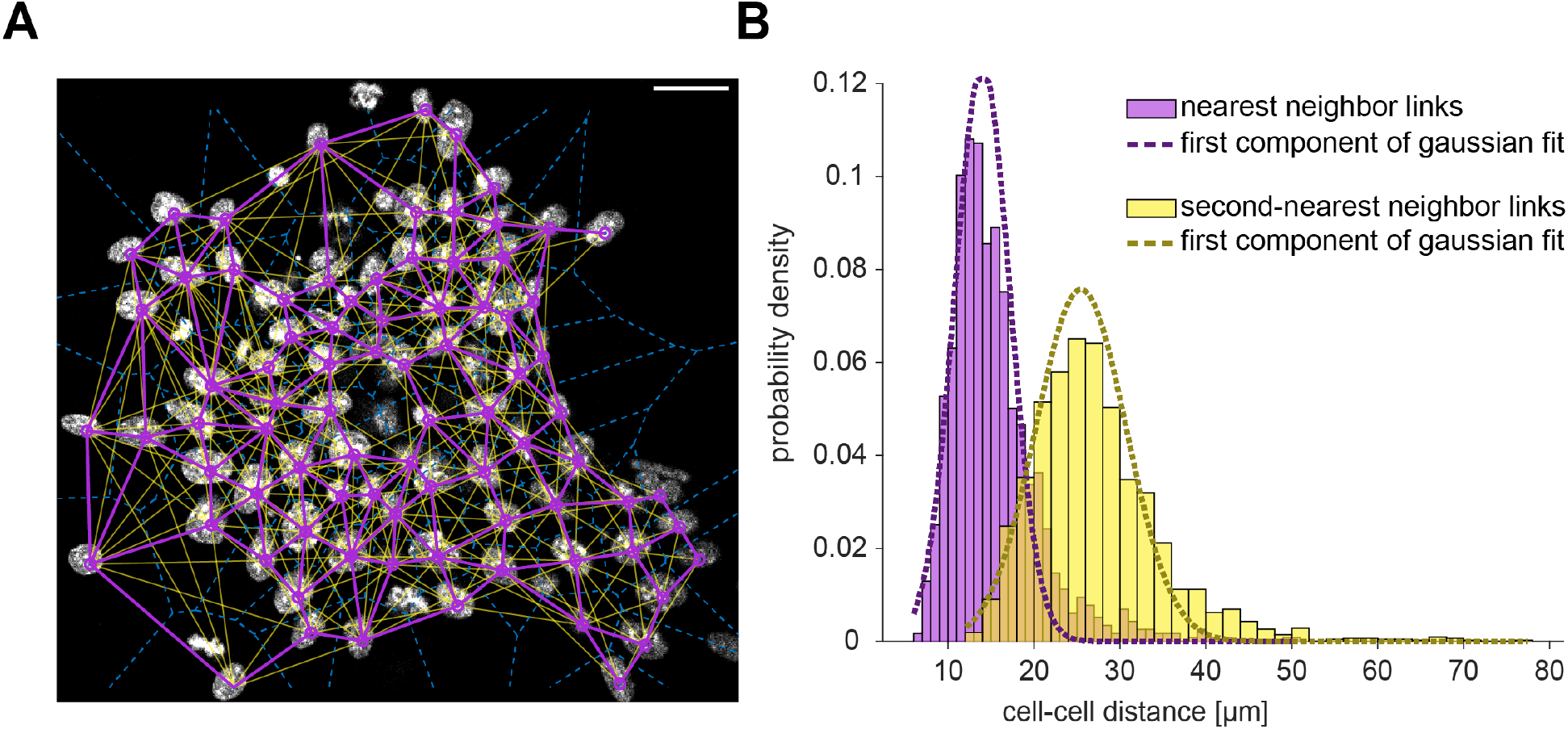
Estimation of cell-cell distances by Delauney triangulation. **A** Nuclear staining (white) of a colony 16 h after initiation of differentiation, overlaid with centers of mass of individual nuclei (purple circles), Voronoi edges (dashed blue lines), and links to nearest (purple) and second nearest neighbors (yellow, see Methods). Scale bar, 20 µm. **B** Histogram of distance distributions between nearest neighbors (purple) and second nearest neighbors (yellow), determined as in **A**. To separate true nearest and second-nearest neighbor distances from spurious longer-distance links, we applied two-component Gaussian mixture model fits to each of the distributions. Dashed lines indicate the first components of each fit. Distances determined by these first components were 14.0 ± 3.2 (mean ± SD) for nearest neighbors, and 25.5 ± 5.3 µm for second nearest neighbors. n = 8 independent fields of view.

**Figure S6.**
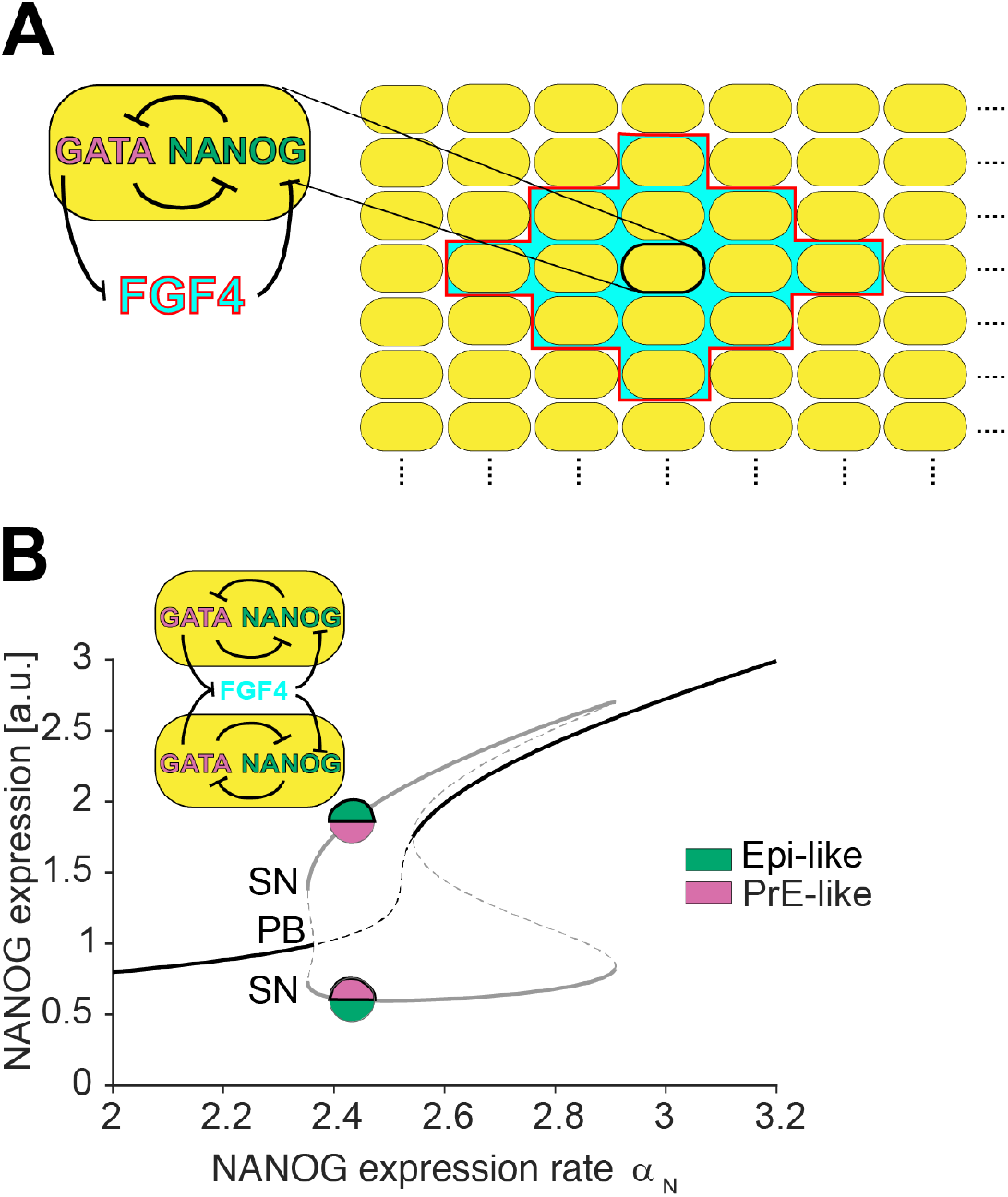
FGF4 signaling range and bifurcation analysis of the model. **A** Scheme illustrating the spatial range of cell-cell communication implemented in the model. FGF4 (cyan with red box) secreted by an individual cell can only reach nearest and second nearest neighbors, and an individual cell will only respond to FGF4 secreted by nearest and second nearest neighbors. **B** Bifurcation diagram showing the principle of emergence of a symmetry-broken inhomogeneous steady state (IHSS) for a two-cell coupled system (inset) in dependence of the production rate constant of NANOG (αN), using NANOG expression as a representative variable. The IHSS (grey lines) is generated via a pitchfork (PB) and stabilized via saddle-node (SN) bifurcations. The two branches represent the heterogeneous attractors: (GATA6-/NANOG+) for cell 1 and (GATA6+/NANOG-) for cell 2, or (GATA6+/NANOG-) for cell 1 and (GATA6-/NANOG+) for cell 2 (representative circles). Solid/dashed lines: stable/unstable solutions. Black: homogeneous steady state; grey: IHSS solution.

**Figure S7.**
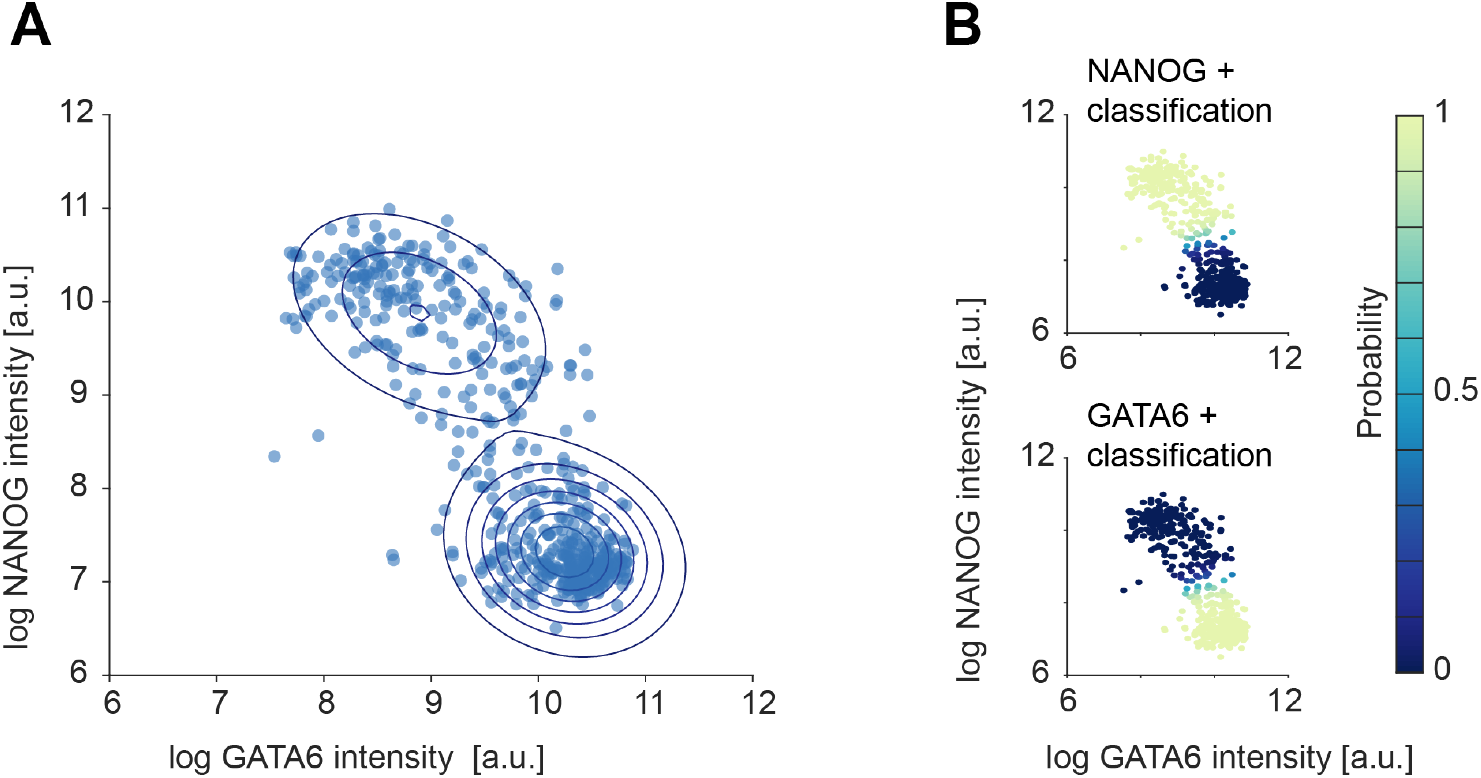
Cell type assignment using a two-component Gaussian mixture model. **A** Blue dots indicate GATA6 and NANOG expression determined by single-cell analysis of immunostaining in cells 16 h after the initiation of differentiation (n=560 cells). Contour lines indicate levels of equal probability density determined from a two-component gaussian mixture model fit. **B** Color-coded posterior probabilities of association with one of the two component distributions of the model fit.

**Figure S8.**
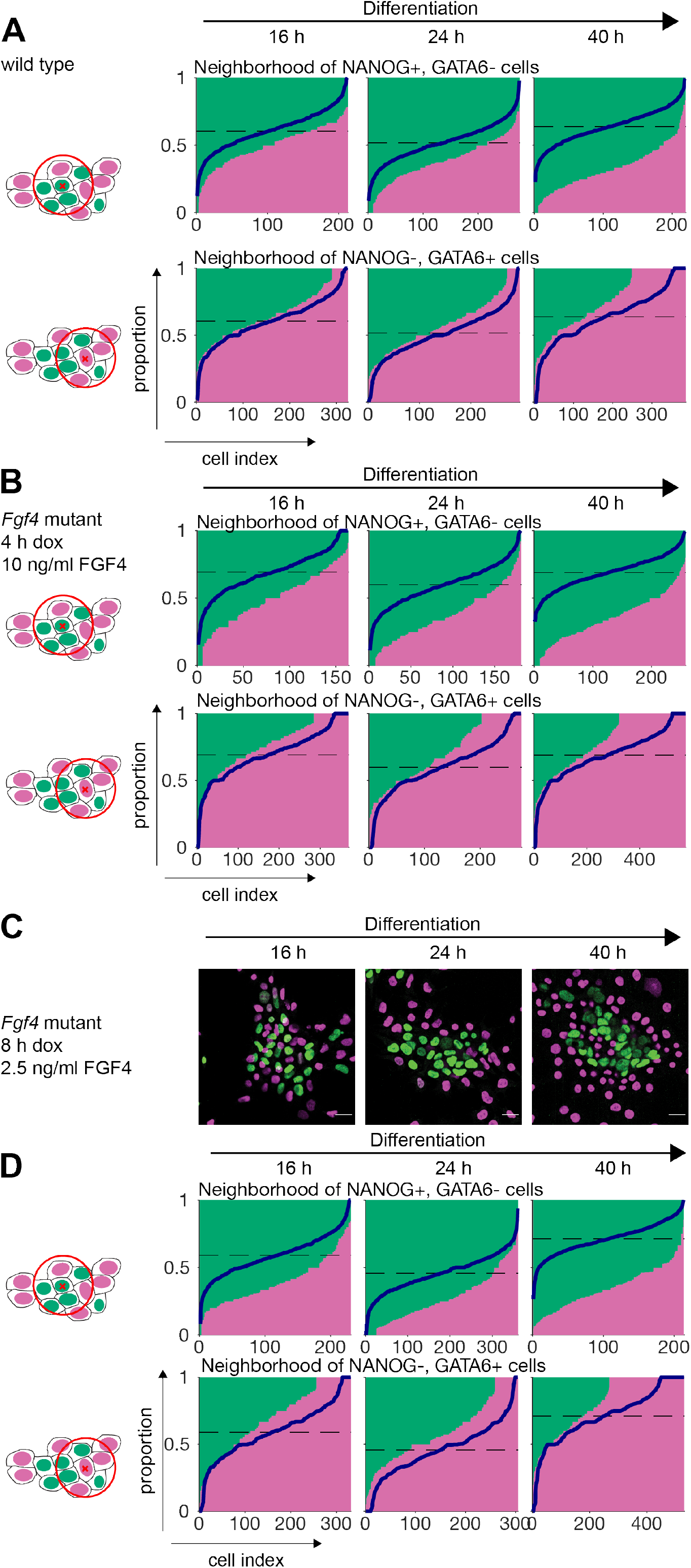
Cell type composition of local cell neighborhoods. **A** Cell type proportions in a local neighborhood of 31.7 µm diameter in wild type cells stimulated for 8 h with doxycycline and differentiated for the indicated times. Each column in a panel corresponds to one cell, and the size of its associated green and magenta bars correspond to the fraction of NANOG+, GATA6- (green) and NANOG-, GATA6+ (magenta) cells in its local neighborhood. Blue lines indicate the corresponding random distribution, calculated by assigning the global distribution of cell types of the time point randomly to experimentally determined cell positions. Clustering of cell types results in the deviation of the true proportions from this random distribution. The local neighborhood of NANOG+, GATA6-cells (top row) indicates clustering of this cell type already at 16 h, which becomes more pronounced over time. Strong clustering of NANOG-, GATA6+ cells (lower row) can only be observed at 40 h. **B** Same analysis as in **A**, but for Fgf4 mutant cells induced for 4 h and differentiated in N2B27 supplemented with 10 ng/ml FGF4 for the indicated periods of time. Deviation from random arrangement (blue lines) indicates clustering of NANOG+, GATA6 cells at 16 h, which becomes more pronounced over time. Some clustering of NANOG-, GATA6+ cells (lower row) can already be detected at 16 h, and becomes more pronounced until 40 h. **C** Immunostaining for NANOG (green) and GATA6 (magenta) in Fgf4 mutant cells induced with doxycycline for 8 h and differentiated in N2B27 supplemented with 2.5 ng/ml FGF4 for the indicated periods of time. **D** Same analysis as in **A, B**, for Fgf4 mutant cells differentiated as described in **C**. Deviations from random arrangement (blue lines) are similar to those seen in **B**, indicating similar clustering dynamics.

**Figure S9.**
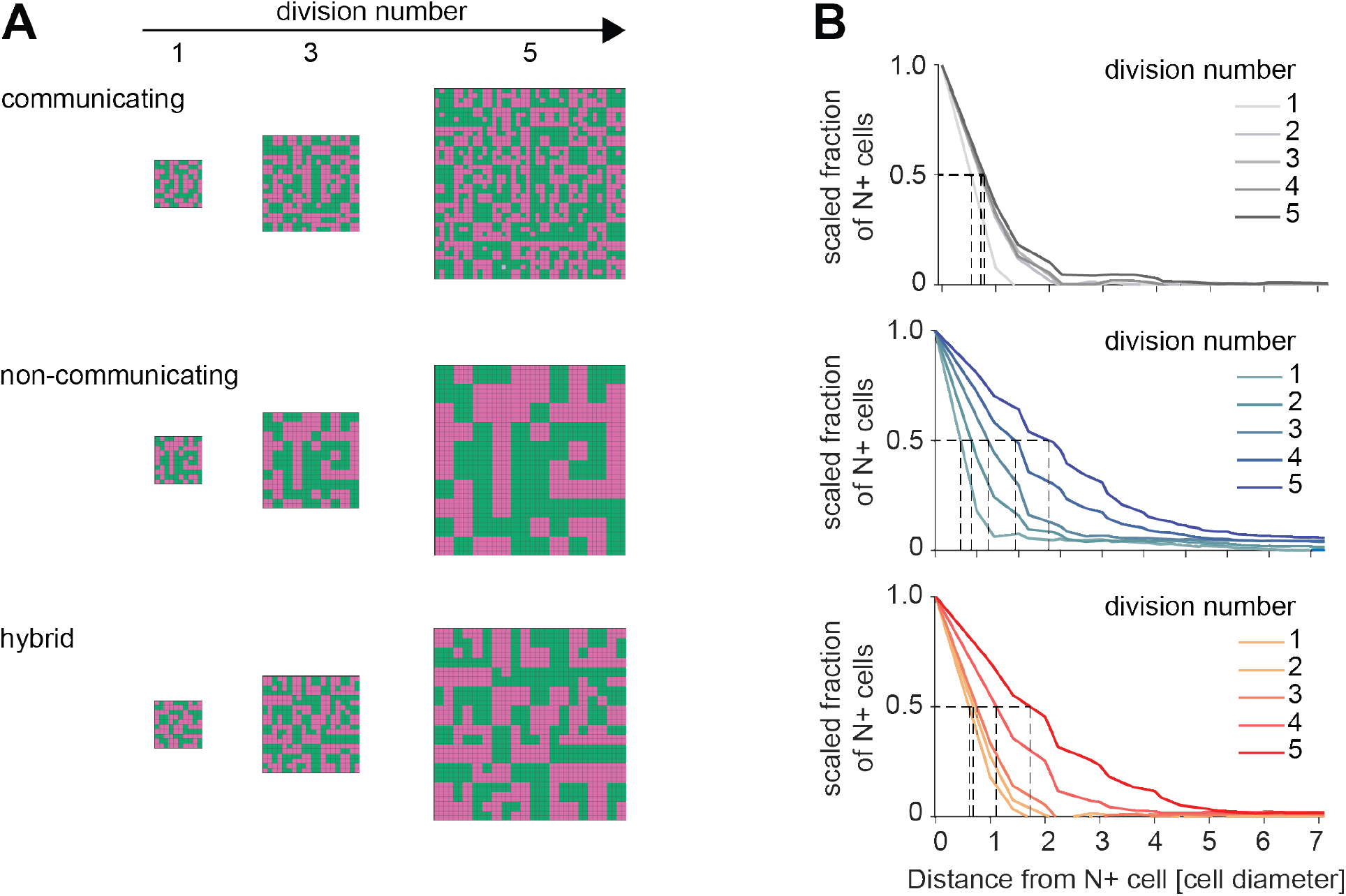
Spatial patterns of cell types in model simulations. **A** Spatial configurations of cell types in single numerical realizations of the model with cell-cell communication (top row), without cell-cell communication (middle row) and of a hybrid model (bottom row) where communication is switched off after the third division. First column: Cell type arrangements at the end of the first cell cycle (grid size 10 x 10), second column - end of the third cell cycle (grid size 20 x 20), third column - end of the fifth cell cycle (grid size 40 x 40). **B** Corresponding estimation of NANOG cluster radius for single numerical realizations of the model variants shown in **A**. Graphs depict the scaled fraction of N+ cells within a specific radius around seed cells. Dashed lines indicate determination of cluster radius, the distance around NANOG+ cells at which the scaled fraction is equal to 0.5. Cluster radii were obtained after each cell cycle of the lineage tree simulation.

**Figure S10.**
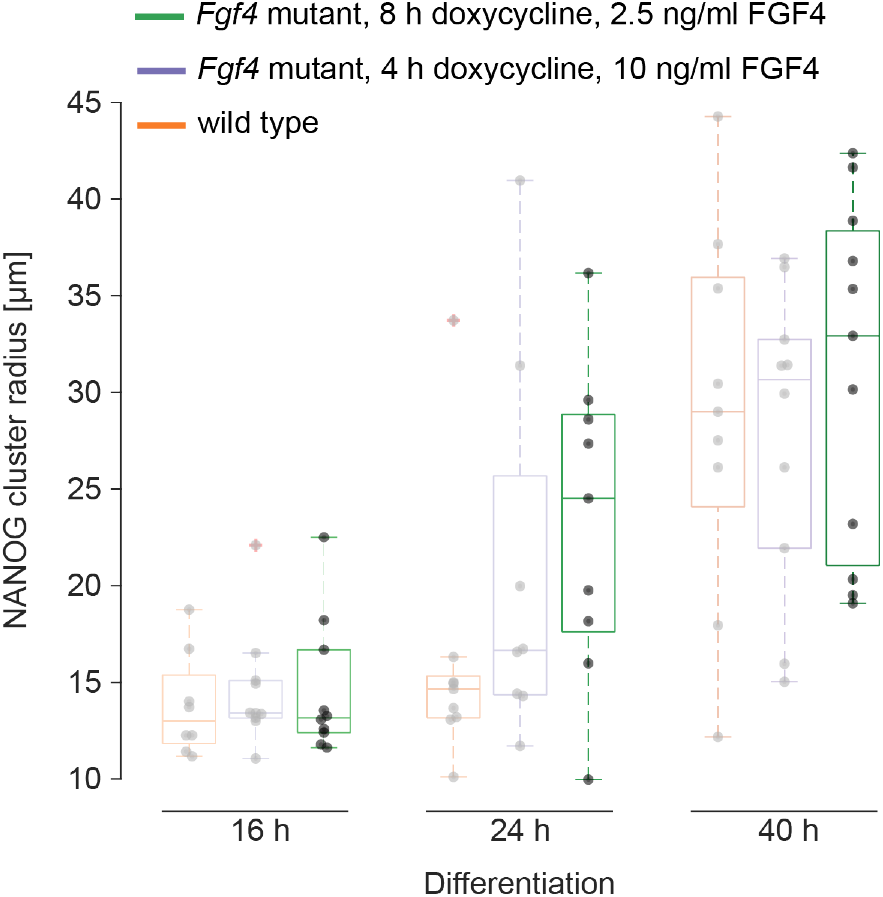
Cluster radius dynamics in Fgf4 mutant cells are independent from stimulation and differentiation regime. Dark: Radius of NANOG clusters in Fgf4 mutant cells induced with doxycycline for 8 h, followed by differentiation in N2B27 medium supplemented with 2.5 ng/ml FGF4 for the indicated times. Dots indicate values from individual fields of view (n ≥ 9), box plots show median, interquartile ranges and outliers (red cross). Shaded data points are reproduced from Figure 4C for comparison, and show the data for Fgf4 wild type cells differentiated in N2B27 only (orange) and for Fgf4 mutant cells differentiated in N2B27 supplemented with 10 ng/ml FGF4 following 4 h of induction with doxycycline.

**Figure S11.**
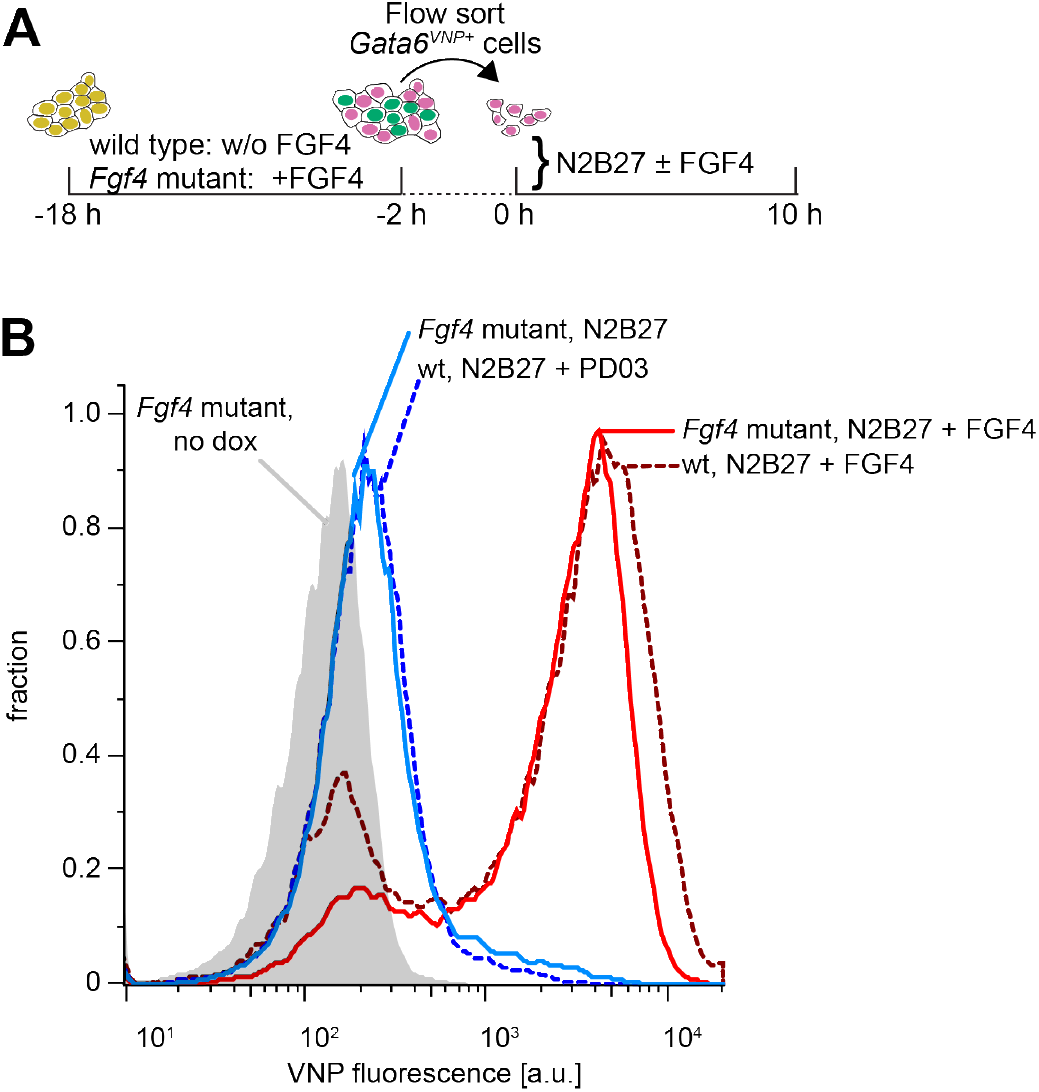
Gata6^VNP^ reporter expression in sorted Fgf4 mutant cells. **A** Schematic representation of the cell-sorting experimental protocol for Fgf4 mutant cells. Following a doxycycline pulse, Fgf4 mutant cells were initially differentiated in medium supplemented with FGF4, while wild type cells were differentiated without recombinant FGF4. After flow sorting of VNP positive cells at 16 h, wild type and Fgf4 mutant cells were cultured in N2B27 medium with or without FGF4, followed by analysis of VNP expression. **B** Flow cytometry histograms of VNP expression in wild type (dashed lines) and Fgf4 mutant (solid lines) inducible Gata6^VNP^ cells, 10 h after sorting of VNP-positive cells and culture in the indicated media.

## Supplementary Movie legends

### Movie S1. Dynamics of VNP expression in unperturbed colonies

Time-lapse imaging of a colony of Gata6^VNP^ reporter cells starting 16 h after the end of a doxycycline pulse. Medium has been switched to N2B27 at the beginning of the recording.

### Movie S2 – S4. Dynamics of VNP expression in sorted cell populations

Time-lapse imaging of Gata6^VNP^ reporter cells flow sorted for VNP expression 16 h after the end of a doxycycline pulse and cultured in defined N2B27 medium alone (Movie S2), or cultured in N2B27 supplemented with 10 ng/ml FGF4 (Movie S3), or cultured in N2B27 supplemented with 1 µM of the MEK inhibitor PD03 (Movie S4).

